# Cell cycle S-phase arrest drives cell extrusion

**DOI:** 10.1101/839845

**Authors:** Vivek K. Dwivedi, Carlos Pardo-Pastor, Rita Droste, Daniel P. Denning, Jody Rosenblatt, H. Robert Horvitz

## Abstract

Cell extrusion is a process of cell elimination in which a cell is squeezed out from its tissue of origin. Extrusion occurs in organisms as diverse as sponges, nematodes, insects, fish and mammals. Defective extrusion is linked to many epithelial disorders, including cancer. Despite broad occurrence, cell-intrinsic triggers of extrusion conserved across phyla are generally unknown. We combined genome-wide genetic screens with live-imaging studies of *C. elegans* embryos and mammalian epithelial cultures and found that S-phase arrest induced extrusion in both. Cells extruded from *C. elegans* embryos exhibited S-phase arrest, and RNAi treatments that specifically prevent S-phase entry or arrest blocked cell extrusion. Pharmacological induction of S-phase arrest was sufficient to promote cell extrusion from a canine epithelial monolayer. Thus, we have discovered an evolutionarily conserved cell-cycle-dependent trigger of cell extrusion. We suggest that S-phase-arrest induced cell extrusion plays a key role in physiology and disease.

## INTRODUCTION

During development and homeostasis, cells are eliminated by a variety of mechanisms. One such mechanism is cell extrusion, in which a cell is expelled from a layer of cells while the continuity of the layer is maintained. Cell extrusion has been observed in and studied using a wide variety of organisms, including the sponge *H. caerulea*, *C. elegans*, *D. melanogaster*, zebrafish and mammals, suggesting that cell extrusion is an evolutionarily conserved mechanism of eliminating unnecessary or harmful cells (De Goeij *et al*., 2009; reviewed by Gudipaty and Rosenblatt, 2017 and Ohsawa *et al*., 2018). Vertebrate epithelial tissues use cell extrusion as the primary mode of cell elimination (Gu and Rosenblatt, 2012; Günther and Seyfert, 2018). Cell extrusion plays a key role in epithelial defense mechanisms that remove oncogene-transformed cells from epithelial layers (reviewed by Kajita and Fujita, 2015). Excessive cell extrusion can produce epithelial layer breaches, like those observed in asthma and Crohn’s disease (Gudipaty and Rosenblatt, 2017). Decreased cell extrusion leads to the formation of epithelial cell masses and confers resistance to cell death (Eisenhoffer *et al*., 2012; Gu *et al*., 2015). Intestinal polyps, which can develop into colon cancers, lack clearly identifiable cell extrusions (Eisenhoffer *et al*., 2012), suggesting that extrusion might be important for the prevention of polyps and intestinal cancers. Disruption and subversion of the cell-extrusion process likely promotes tumor growth and metastasis in pancreatic, lung and colon cancer (Gu *et al.*, 2015).

While several mechanisms of cell extrusion have been described for *Drosophila* and vertebrates, these mechanisms have focused on the cell-cell interactions and cellular contexts, such as crowding, topological defects, cell competition, etc., that induce cell extrusion (reviewed by Fadul and Rosenblatt, 2018, Ohsawa *et al*., 2018). A cell-intrinsic trigger that can induce extrusion in organisms of different phyla has not been identified.

*C. elegans* is an excellent organism for the study of evolutionarily conserved mechanisms of cell elimination. The discovery of a conserved set of genes regulating caspase-mediated apoptosis in *C. elegans* has been fundamental to the understanding of programmed cell death in metazoa (reviewed by Fuchs and Steller, 2011). Cell extrusion can eliminate cells fated for death in *C. elegans* (Denning *et al*., 2012). Embryos with mutations in the caspase-mediated apoptosis pathway, e.g. loss-of-function mutants of the caspase gene *ced-3*, eliminate by extrusion a subset of cells that are otherwise eliminated by caspase-mediated apoptosis and engulfment in wild-type embryos. Denning *et al*. (2012) determined that the PAR-4 – PIG-1 (mammalian homologs LKB1 – MELK) kinase cascade is required for cell extrusion by *C. elegans ced-3(lf)* embryos. However, LKB1 (mammalian homolog of PAR-4) was found to prevent extrusion from mouse embryos (Krawchuk *et al*., 2015), indicating that LKB1/PAR-4 is likely not a driver but a regulator of cell extrusion in nematodes and mammals. No “caspase-equivalent” cell-intrinsic driver of cell extrusion is known.

To seek a conserved cell-intrinsic driver of extrusion, we first comprehensively identified genes and pathways that control cell extrusion by *C. elegans* and then tested the corresponding pathways for a role in mammalian cell extrusion. Briefly, we performed a genome-wide RNAi screen for defective cell extrusion by *C. elegans* and used confocal microscopy to analyze the effect of RNAi against the identified genes on the cell extrusion process. From this analysis, we found that cell extrusion by *C. elegans* requires cell-autonomous cell cycle entry and subsequent S-phase arrest and that circumventing S-phase arrest blocks extrusion. We then tested and confirmed that pharmacological induction of S-phase arrest with hydroxyurea (HU) (Timson, 1975; Bianchi *et al*., 1983) promotes cell extrusion of mammalian epithelial cells. We conclude that S-phase arrest is a conserved cell-intrinsic trigger of cell extrusion in *C. elegans* and mammals.

## RESULTS

### Genome-wide RNAi screen identified cell-cycle genes as candidate regulators of cell extrusion

In wild-type *C. elegans* embryos, 131 cells are eliminated by caspase-mediated apoptosis and engulfment. By contrast, in *C. elegans ced-3* caspase mutants, a few of the cells that would normally undergo programmed cell death instead are extruded from the developing embryo (Denning *et al*., 2012). Of the approximately six cells extruded from *ced-3(lf)* embryos, the cell ABplpappap is most frequently extruded (Denning *et al*., 2012 and unpublished). If extrusion fails to occur, ABplpappap (or descendants of ABplpappap) survive(s) and differentiate(s) into one (or two) supernumerary excretory cell(s), producing mutant animals with the two (or three)-excretory-cell (Tex) phenotype; by contrast both wild-type and *ced-3(lf)* animals have one excretory cell (Denning *et al*., 2012; Figure 1A). Mutations that reduce ABplpappap extrusion and produce the Tex phenotype also reduce the extrusion of other cells (Denning *et al*., 2012), making the Tex phenotype a convenient marker for defective cell extrusion.

**Figure 1.**
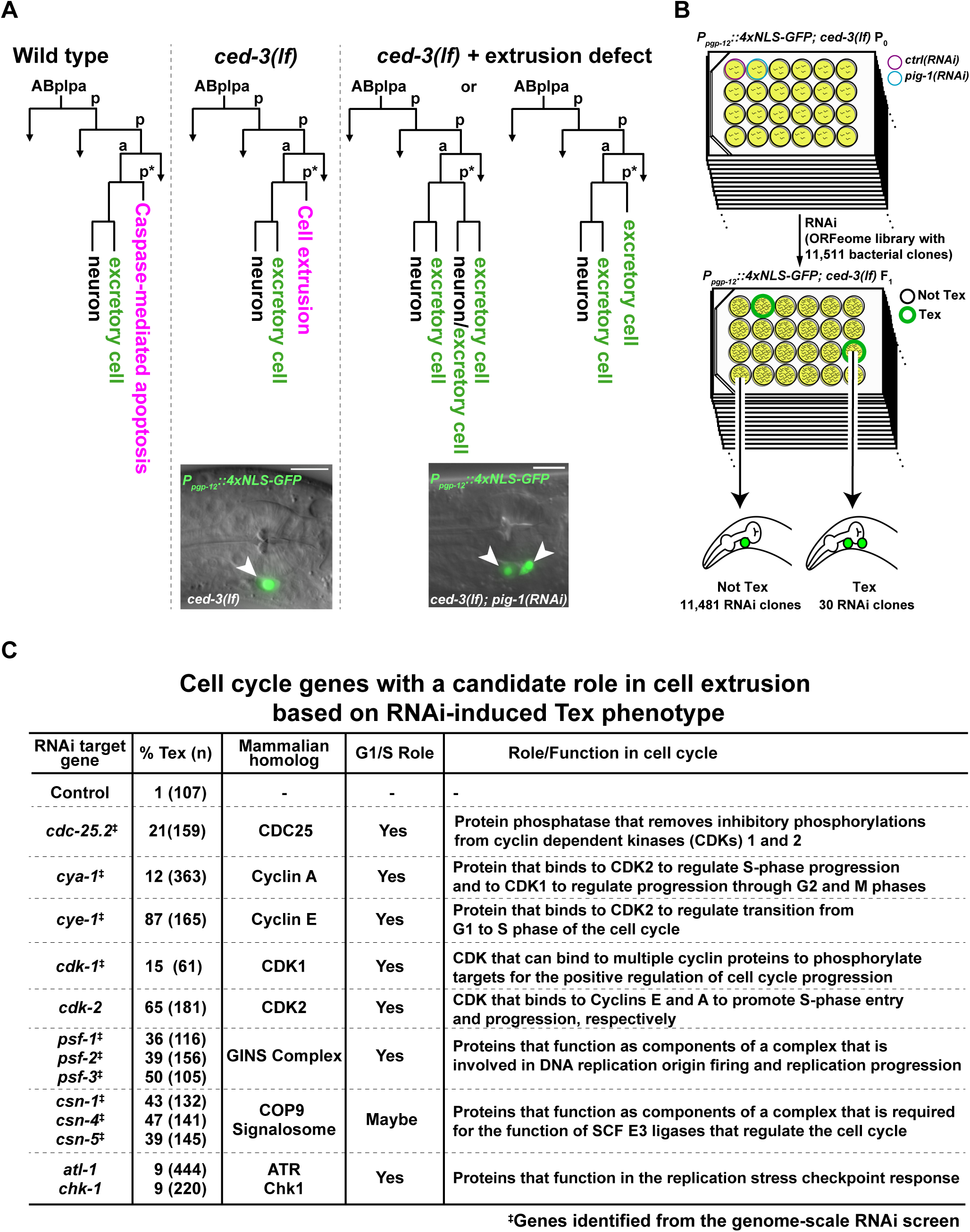
A Genome-wide RNAi screen for genes required for cell extrusion identifies multiple cell-cycle genes. (A) Lineage diagrams show the fate of the cell ABplpappap (marked with an *) in embryos that are wild-type, embryos with a *ced-3(lf)* mutation and embryos with a *ced-3(lf)* mutation and a defect in cell extrusion. A micrograph of the pharyngeal region showing excretory cell(s), which express nuclear GFP and are marked with white arrowhead(s), is shown for a representative *ced-3(lf)* animal and a *ced-3(lf)* + extrusion defect (*ced-3(lf); pig-1(RNAi)*) animal below the corresponding cell-lineage diagrams. Scale bar, 10 µm. (B) Schematic representation of the genome-wide RNAi screen for the Tex phenotype. (C) RNAi clone targets with a function in the cell cycle identified from the genome-wide or candidate-based RNAi screens for the Tex phenotype and the corresponding penetrance of the Tex phenotype. The mammalian homologs of these genes and their functions in mammals are shown. G1/S Role indicates whether an identified RNAi target has a role in G1, G1-to-S phase transition or S-phase progression in mammals. ‡, genes identified from the genome-wide RNAi screen; other genes were identified from a candidate RNAi screen of cell cycle genes for the Tex phenotype (Supplemental Table 1).

We performed a genome-wide RNAi screen for the Tex phenotype in *ced-3(lf)* animals expressing the GFP excretory-cell reporter *P_pgp-12_::4xNLS::GFP* (Figure 1B; Denning *et al*., 2012). We screened 11,511 RNAi clones (targeting about 55% of the ∼20,000 *C. elegans* genes by feeding (Rual *et al*., 2004)) and found 30 clones targeting 27 unique genes that consistently produced a Tex phenotype. Three RNAi clones identified genes previously reported to function in cell extrusion, *grp-1*, *arf-1.2* and *arf-3* (Denning *et al*., 2012), confirming that this RNAi screen could identify cell extrusion mutants. Unexpectedly, 10 of the RNAi clones targeted genes that control cell-cycle progression (Figure 1C), suggesting a possible role for the cell cycle in controlling cell extrusion.

We then tested a nearly complete set of *C. elegans* cell cycle genes for functional roles in cell extrusion (van den Heuvel, 2005) using an RNAi library of 61 publically available and, when necessary, newly generated clones with each clone targeting a unique gene (Kamath *et al*., 2003; Rual *et al.*, 2004; Materials and Methods). We found that RNAi against four additional cell cycle genes produced a Tex phenotype (Figure 1C, Supplemental Table 1).

Most of the 14 cell-cycle genes we identified are well-characterized regulators of S-phase entry and progression. These genes include *cdc-25.2*, which encodes a homolog of the CDK-activating phosphatase CDC25 (Lee *et al*., 2016); *cdk-1* and *cdk-2*, which encode homologs of CDK1 and CDK2, respectively (Boxem, 2006); *cye-1* and *cya-1*, which encode homologs of S-phase cyclins E and A, respectively (Fay and Han, 2000; Kreutzer *et al*., 1995); and *psf-1*, *psf-2* and *psf-3*, which encode homologs of pre-replicative and replicative complex components PSF1, PSF2 and PSF3, respectively (Ossareh-Nazari *et al*., 2016; Figure 1C). All of these cell cycle genes with S-phase function are required for *C. elegans* viability. However, it is unlikely that a general reduction in embryonic fitness causes the Tex phenotype, as RNAi against essential genes involved in other phases of the cell cycle (e.g., metaphase-to-anaphase transition genes *mat-1*, *mat-2*, etc. (reviewed by Yeong, 2004)) or those involved in transcription (e.g., *cdk-7* and *cdk-9* (Wallenfang and Seydoux, 2002; Bowman *et al*., 2013)) did not produce the Tex phenotype despite producing extensive lethality (Supplemental Table 1). Furthermore, as with RNAi targeting *pig-1* or other genes generally required for cell extrusion (Denning *et al*., 2012), RNAi against 13 of the 14 identified cell cycle genes produced the Tex phenotype in *ced-3(lf)* mutants but not in wild-type animals (Supplemental Table 2). Altogether, we found that defects in S-phase entry and progression are specifically associated with a synthetic *ced-3*-dependent Tex phenotype, suggesting a role for these cell-cycle genes in promoting cell extrusion.

### S-phase entry genes function to promote cell extrusion

Whereas defects in the extrusion of ABplpappap cause the Tex phenotype, it is possible that the supernumerary excretory cell(s) of some mutants could arise from other cell lineages. To directly determine whether genes functioning in S-phase entry are important for cell extrusion, we used time-lapse confocal microscopy to monitor the extrusion of ABplpappap from *ced-3(lf)* embryos deficient in *cye-1* or *cdk-2*. To assess extrusion events, we imaged live embryos with ABplpappap in focus over a 50-minute period ending at the completion of ventral enclosure, when epidermal cells meet at the ventral midline following a dorsolateral migration; ventral enclosure coincides with the cell extrusion that occurs in *ced-3(lf)* embryos (Denning *et al*., 2012). For clarity, we refer below to embryos from *ced-3(lf)* parents treated with RNAi against a gene, say *gene-x*, as “*gene-x(RNAi)* embryos” and to embryos from *ced-3(lf)* parents treated with RNAi against the empty vector as “control embryos.”

In control embryos, cells neighboring ABplpappap on the ventral surface gradually disappeared from view until ABplpappap was left completely isolated (Figure 2A, Movie 1), indicating that ABplpappap had been extruded from the embryo. By comparison, in *cye-1(RNAi)* embryos (Figure 2B, Movie 2) or *cdk-2(RNAi)* embryos (Figure 2C, Movie 3), ABplpappap was surrounded by cells throughout the imaging period and hence remained within the embryo, failing to detach. Using dorso-ventral confocal sections from the live embryo imaging, we reconstructed sagittal views of the embryos during ventral enclosure. These views confirmed that ABplpappap was extruded ventrally from 10 of 11 control embryos (Figure 2D, Supplemental Figure 1), whereas ABplpappap failed to detach and was incorporated into the body of *cye-1(RNAi)* embryos (11 of 11 embryos; Figure 2E, Supplemental Figure 2) or *cdk-2(RNAi)* embryos (10 of 11 embryos; Figure 2F, Supplemental Figure 3). These findings demonstrate that the S-phase entry genes *cye-1* and *cdk-2* are required for ABplpappap extrusion in *ced-3* embryos.

**Figure 2.**
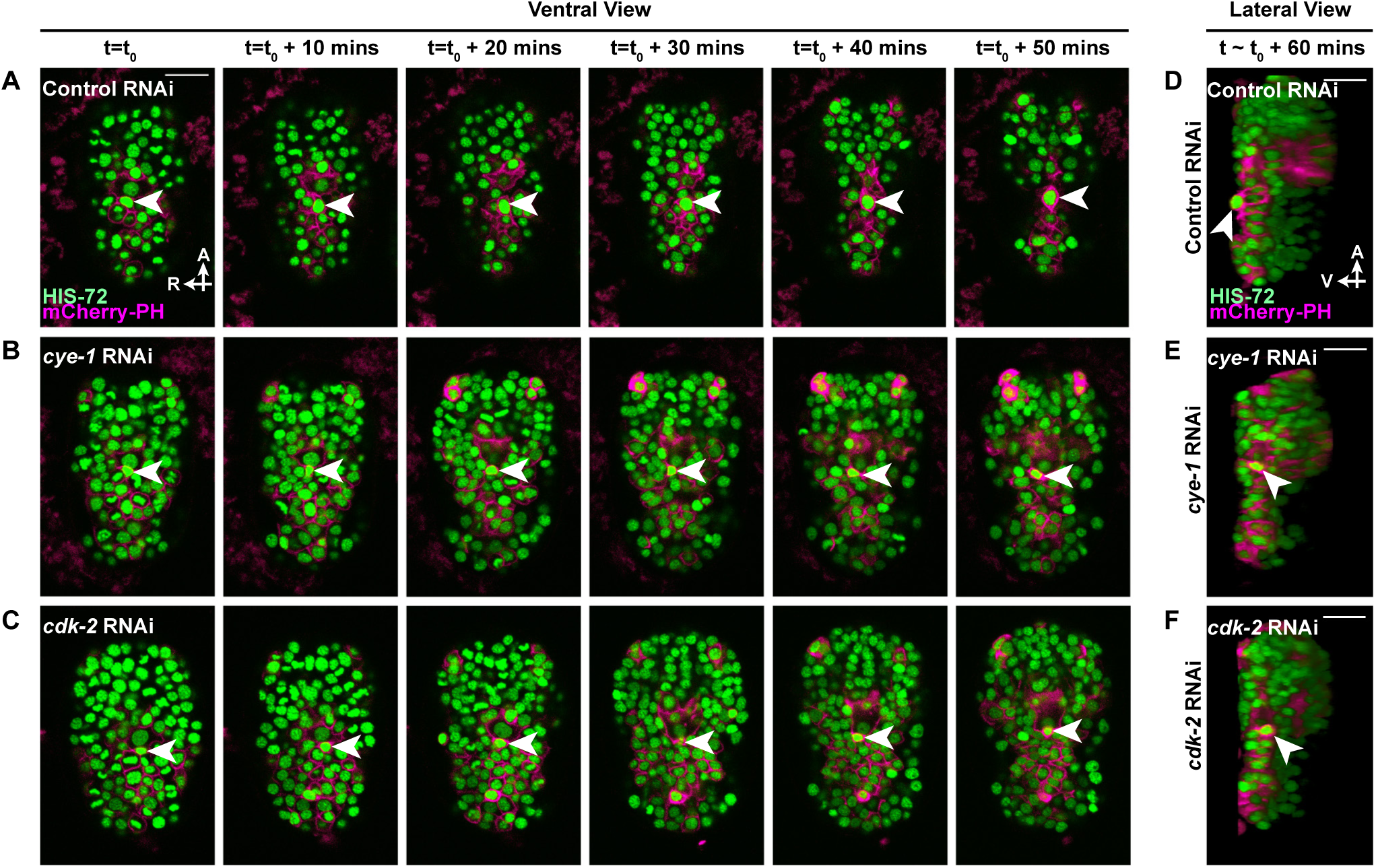
Cell extrusion of ABplpappap requires the function of the cell-cycle genes *cye-1* and *cdk-2*. (A-C) Micrographs of the ventral surface obtained at 10-min intervals over a period of 50 min using time-lapse confocal microscopy show the location of ABplpappap (arrowhead) on the ventral surface with respect to other embryonic cells in (A) control, (B) *cye-1(RNAi)*, and (C) *cdk-2(RNAi)* embryos. The embryos shown carried the transgenes *stIs10026* and *nIs632*. A, anterior; R, right. Scale bar, 10 µm. (D-F) Virtual lateral sections through the ABplpappap cell in *ced-3(lf)* embryos show the relative location of ABplpappap (arrowhead) in (D) control, (E) *cye-1(RNAi)*, and (F) *cdk-2(RNAi)* embryos. Embryos shown in (D), (E) and (F) are the same as those in (A), (B) and (C), respectively. A, anterior; V, ventral. Scale bar, 10 µm.

### Entry into S phase is required for and precedes cell extrusion

Since genes that promote S-phase entry are required for ABplpappap extrusion, we tested if cells that undergo extrusion enter S phase. We used a previously characterized reporter transgene that expresses a truncated human DNA Helicase B (tDHB)-GFP fusion protein optimized for expression in *C. elegans* and that changes its intracellular location in response to CDK1 and CDK2 activity (van Rijnberk *et al*., 2017; Spencer *et al*., 2013). tDHB-GFP is enriched in the nuclei of quiescent or post-mitotic cells, whereas it exhibits an increasing cytoplasmic bias as cells progress from S-phase through mitosis (Figure 3A; Spencer *et al*., 2013). In control embryos, tDHB-GFP was mostly absent from the ABplpappap nucleus (10 of 10 embryos) both before ventral enclosure (Figure 3B) and as it was extruded (Figure 3E), indicating that ABplpappap entered S phase prior to its extrusion during the period of ventral enclosure. Cells extruded from other sites of the embryo also displayed low levels of nuclear tDHB-GFP (Figures 3I-L). By contrast, in *cye-1(RNAi)* embryos (Figures 3C, 3F) or *cdk-2(RNAi)* embryos (Figures 3D, 3G) the ABplpappap nucleus scored positive for the tDHB-GFP fusion protein during the period around ventral enclosure (10 of 10 embryos each for each RNAi treatment), suggesting that RNAi against these genes prevented both the entry of ABplpappap into S-phase and its extrusion. Quantification of the nuclear-to-cytoplasmic ratio of tDHB-GFP fluorescence intensity in ABplpappap in control, *cye-1(RNAi)* and *cdk-2(RNAi)* embryos at varying stages with respect to ventral enclosure confirmed these observations (Figure 3H). Some cells were still extruded in *cye-1(RNAi)* and *cdk-2(RNAi)* embryos, likely reflecting incomplete inhibition of gene function by RNAi. Consistently, such cells displayed low levels of nuclear tDHB-GFP, indicating that those cells entered the cell cycle (Supplemental Figure 4). Thus, cell-cycle entry appears to be a functionally critical step in the process of cell extrusion.

**Figure 3.**
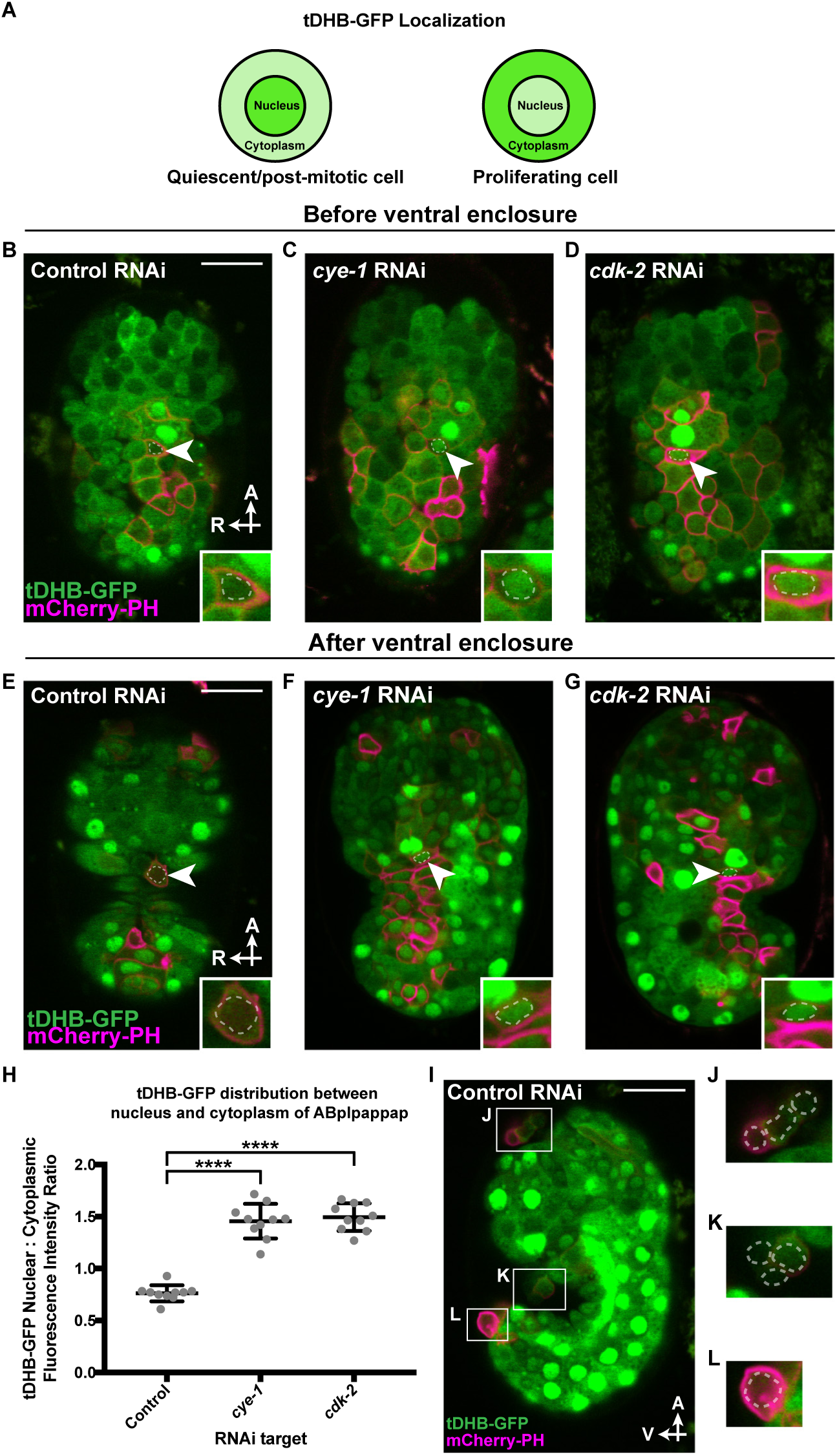
Cells that are extruded enter the cell cycle and are extruded in a *cye-1*- and *cdk-2* -dependent manner. (A) Schematic showing the relative nuclear/cytoplasmic localization of a truncated DHB (tDHB) – GFP fusion protein in a quiescent/post-mitotic cells and in cells between S phase and mitosis in the cell cycle (van Rijnberk *et al*., 2017). (B-G) Micrographs of the ventral surface, including ABplpappap (arrowhead), of *ced-3(lf)* embryos expressing tDHB-GFP obtained (B-D) prior to and (E-G) post ventral enclosure using confocal microscopy in (B,E) control embryos, (C,F) *cye-1 (RNAi)* embryos, and (D,G) *cdk-2(RNAi)* embryos. Inset, a magnified view of the ABplpappap cell. The ABplpappap nucleus, as determined by Nomarski optics, is marked by a dotted line in each image and inset. The embryos shown carried transgenes *heSi192* and *nIs861*. A, anterior; R, right. Scale bar, 10 µm. (H) Quantification of the ratio of tDHB-GFP fluorescence intensity in the nucleus to that in the cytoplasm in control, *cye-1(RNAi)*, or *cdk-2(RNAi)* embryos expressing tDHB-GFP. ****, p<0.0001 per ordinary one-way ANOVA of the log of ratios. n=10 embryos per RNAi treatment. (I-L) Micrograph of (I) a *ced-3(lf)* embryo expressing tDHB-GFP obtained at the comma stage using confocal microscopy. Magnified views are provided for the cells extruded at (J) the anterior sensory depression, (K) the ventral pocket, and (L) the posterior tip of the embryo, with the corresponding regions outlined in (I). The cell nucleus, as determined by Nomarski optics, is marked by a dotted line for each extruded cell in the magnified images (J), (K) and (L). The embryo shown carried the transgenes *heSi192* and *nIs861*. A, anterior; V, ventral. Scale bar, 10 µm.

To define more precisely the cell-cycle phase that facilitates cell extrusion, we used a second reporter transgene (GFP::PCN-1), which expresses an N-terminal translational fusion of GFP to the *C. elegans* homolog of the DNA replication processivity factor PCNA (Brauchle *et al*., 2003). PCNA in mammalian cells and early *C. elegans* embryonic cells exhibits a punctate, sub-nuclear localization only during S-phase (Figure 4A; Brauchle *et al*., 2003; Zerjatke *et al*., 2017). The localization pattern of GFP::PCN-1 in cell cycles of cells close to ABplpappap on the embryonic ventral surface matched that described for early embryonic cells and contrasted with the continuous accumulation of GFP::PCN-1 observed during the *C. elegans* germline cell cycle (Supplemental Figure 5; Movie 4; Brauchle *et al*., 2003; Kocsisova *et al*., 2018). We found that in control embryos GFP::PCN-1 was localized in bright sub-nuclear foci in ABplpappap immediately prior to the initiation of ventral enclosure, indicating that this cell was in S phase (5 of 5 embryos) (Figure 4B). By contrast, ABplpappap showed a diffuse nuclear localization of GFP::PCN-1 in *cye-1(RNAi)* (Figure 4C, 4F) and *cdk-2(RNAi)* (Figure 4D, 4G) embryos, both before (5 of 5 embryos each) and after ventral enclosure (5 of 5 embryos each), indicating a failure to enter the cell cycle. We conclude that cells to be extruded must enter S phase for extrusion to occur and that blocking S-phase entry prevents cell extrusion.

**Figure 4.**
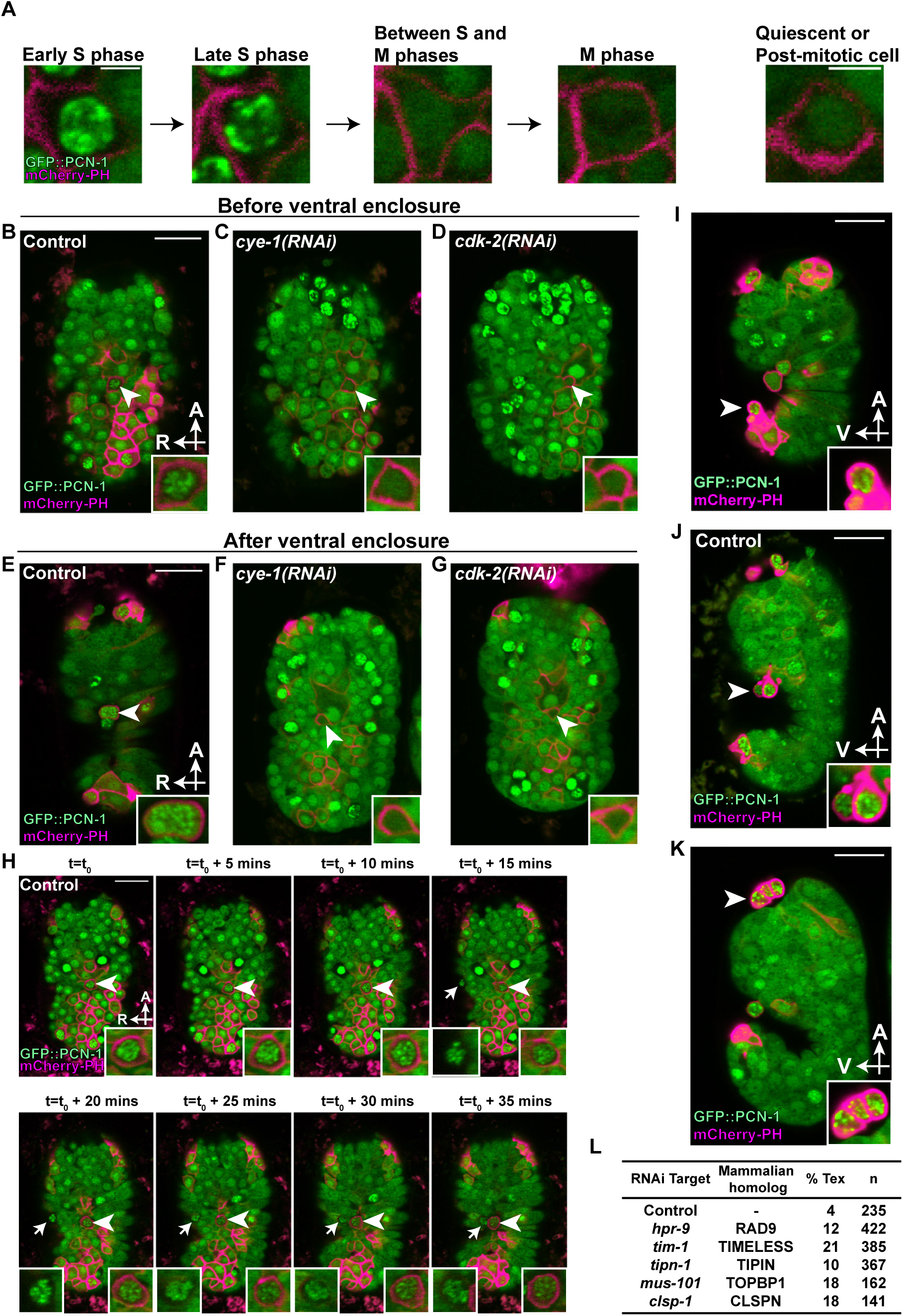
Cells undergoing extrusion arrest in S phase and trigger the DNA replication stress checkpoint. (A) Representative micrographs showing localization patterns of GFP::PCN-1 in the same cell at different phases of the cell cycle or in a different cell in a post-mitotic state. The post-mitotic cell shown is ABplpappaaa (future RMEV neuron). These cells were imaged from a *ced-3(lf)* embryo, which carried the transgenes *nIs861* and *isIs17*. Scale bar, 2 µm. (B-G) Micrographs of the ventral surface, including ABplpappap (arrowhead), of *ced-3(lf)* embryos expressing GFP::PCN-1 obtained (B-D) prior to and (E-G) post ventral enclosure using confocal microscopy in (B,E) control embryos, (C,F) *cye-1(RNAi)* embryos, and (D,G) *cdk-2(RNAi)* embryos. Inset, a magnified view of ABplpappap. The embryos shown carried the transgenes *nIs861* and *isIs17.* A, anterior; R, right. Scale bar, 10 µm. (H) Micrographs obtained at 5-min intervals using time-lapse confocal microscopy show the GFP::PCN-1 localization pattern in ABplpappap (arrowhead) and another unidentified extruded cell (arrow) as they were progressively extruded over 35 min in a *ced-3(lf)* embryo with control RNAi. Left insets, magnified views of unidentified extruded cell; right insets, magnified views of ABplpappap. The embryo shown carried the transgenes *isIs17* and *nIs861*. The decrease in fluorescence intensity can be attributed to bleaching from repeated imaging over time. A, anterior; R, right. Scale bar, 10 µm. (I-K) Micrographs of extruded cells in *ced-3(lf)* embryos expressing GFP::PCN-1 obtained at the comma stage using confocal microscopy show GFP::PCN-1 localization in cells at (I) the posterior tip of the embryo (no RNAi), (J) the ventral pocket (control RNAi), and (K) the anterior sensory depression (no RNAi). Inset, a magnified view of extruded cells marked by an arrowhead. The embryos shown carried the transgenes *isIs17* and *nIs861*. A, anterior; V, ventral. Scale bar, 10 µm. (L) Penetrances of the Tex phenotype produced by RNAi-mediated targeting of replication-stress checkpoint genes *hpr-9*, *tim-1*, *tipn-1*, *mus-101* and *clsp-1* in *ced-3(lf)* animals. The strain used for scoring the RNAi-induced Tex phenotype carried the transgene *nIs433*, which expresses nuclear GFP in excretory cell(s).

### Extruding cells exhibit an arrested S phase and experience replication stress

Given that cells required cell-cycle entry for extrusion, we examined the extent of cell-cycle progression in these cells as they were extruded. In control embryos, we found that GFP::PCN-1 was localized in bright sub-nuclear foci in ABplpappap both before (5 of 5 embryos) and after extrusion (5 of 5 embryos) (Figure 4B, 4E), indicating that ABplpappap entered but did not exit S phase. We observed no significant changes of GFP::PCN-1 localization in ABplpappap up to and after its extrusion over a period of 35 min (Figure 4H), indicating that it remained arrested in S phase. A second unidentified extruding cell showed a similarly unchanging GFP::PCN-1 localization pattern (Figure 4H). To determine if the S-phase arrest observed in ABplpappap and the other extruding cell is a general feature of cell extrusion, we examined other extruded cells in the embryo. Nearly all cells extruded by *ced-3(lf)* embryos displayed bright sub-nuclear foci of GFP::PCN-1, consistent with an arrested S phase in these cells (Figures 4H-J).

Arrest of DNA replication during S phase can occur as a result of replication stress, which can arise for many different reasons (reviewed by Zeman and Cimprich, 2014). Replication stress triggers the replication stress response, which stabilizes stalled replication forks, halts cell-cycle progression and prevents further firing of replication origins (reviewed by Zeman and Cimprich, 2014). As cells undergoing extrusion are arrested in S-phase, we asked if triggering the replication stress response was important for extrusion. Core components of the replication stress response pathway in *C. elegans* and other metazoans include ATR, Chk1, Rad17, Rad9, Rad1, Hus1, Replication Protein A, TopBP1, Timeless, Tipin and Claspin proteins (Stevens *et al*., 2016; Yazinski and Zou, 2016). RNAi against 7 of the 11 *C. elegans* genes encoding these core components of the replication-stress checkpoint (*atl-1*, *chk-1*, *hpr-9*, *mus-101*, *tim-1*, *tipn-1* and *clsp-1*) produced a Tex phenotype in *ced-3(lf)* animals (Figure 1C, Figure 4L), indicating an involvement of replication stress response in cell extrusion. These findings suggest that the replication stresses underlying the S-phase arrest in extruding cells trigger the replication stress response, which promotes cell extrusion.

### Bypassing S-phase arrest and completing the cell cycle prevents cell extrusion

After determining that S-phase arrest is a key feature of cell extrusion, we asked whether the previously identified *C. elegans* cell extrusion regulators *pig-1* (homolog of the mammalian kinase gene *MELK*) and *grp-1* (homolog of the mammalian ARF GEF gene *CYTH3*) (Denning *et al*., 2012) also play a role in producing this S-phase arrest. We monitored the fate of ABplpappap in *pig-1(RNAi)* embryos by time-lapse confocal microscopy. Strikingly, we found that instead of undergoing S-phase arrest, ABplpappap completed the cell cycle and divided before ventral enclosure in these embryos (Figure 5A; Movie 5). By examining virtual lateral sections, we found that instead of undergoing extrusion, as in the case of control embryos (Figure 5B), ABplpappap in *pig-1(RNAi)* embryos divided to generate daughters that were not extruded (5 of 6 embryos) (Figure 5C). The same fate of ABplpappap was observed in *grp-1(RNAi)* embryos (5 of 5 embryos) (Figure 5D). These findings indicate that in addition to the genes *cye-1* and *cdk-2*, the genes *pig-1* and *grp-1* are required to produce the S-phase arrest that precedes cell extrusion.

**Figure 5.**
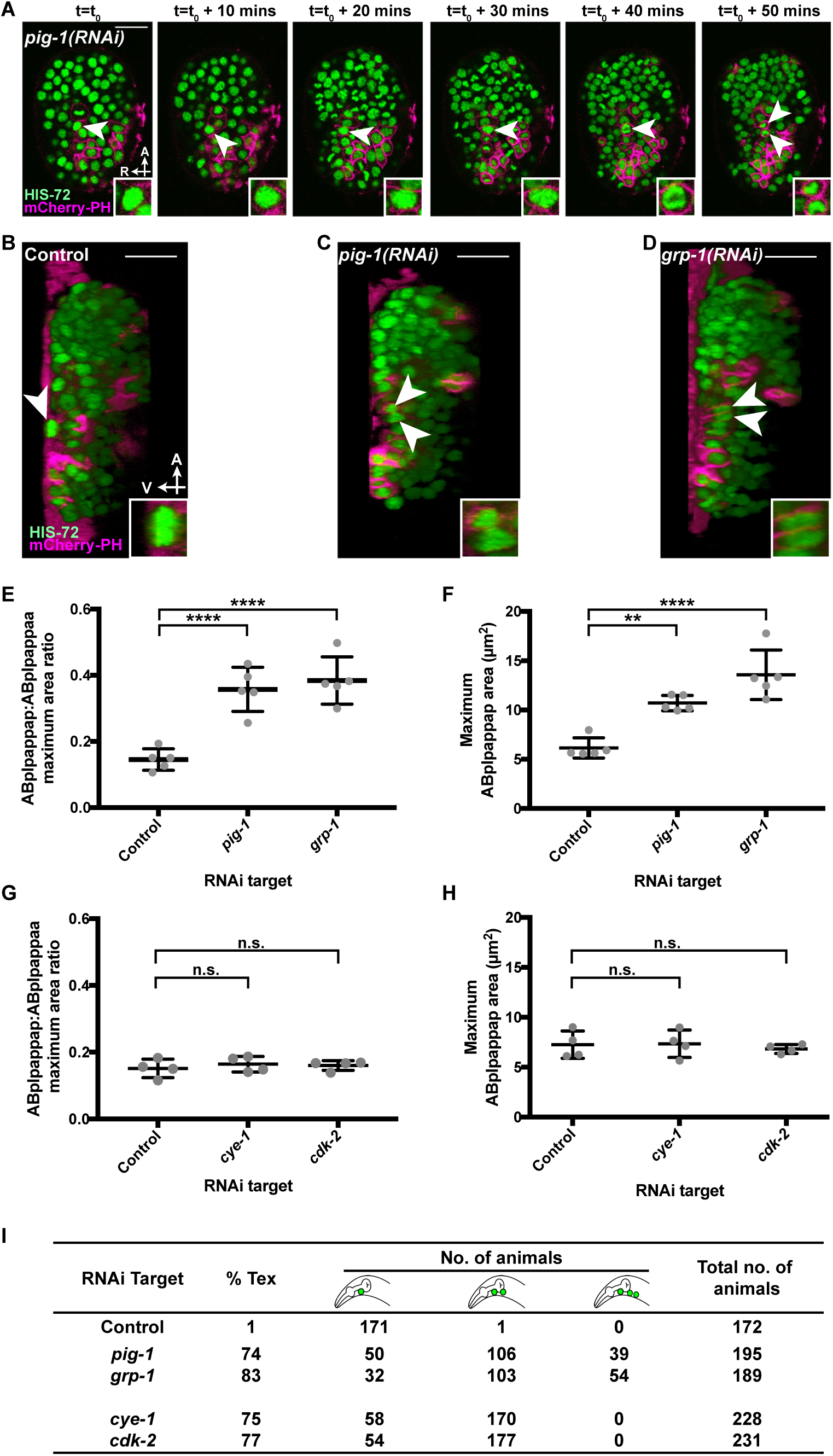
Unequal cell division genes *pig-1* and *grp-1* promote extrusion by preventing cell-cycle progression beyond S phase. (A) Micrographs of a *pig-1(RNAi)* embryo obtained at 10-min intervals over a period of 50 min using time-lapse confocal microscopy show that an enlarged ABplpappap divides before ventral enclosure in *pig-1(RNAi)* embryos. The embryo shown carried the transgenes *stIs10026* and *nIs861*. Inset, a magnified view of ABplpappap or its descendants. A, anterior; R, right. Scale bar, 10 µm. (B-D) Virtual lateral sections of *ced-3(lf)* embryos through ABplpappap (single arrowhead) or its daughters (two arrowheads) in a (B) control embryo, (C) *pig-1(RNAi)* embryo, and (D) *grp-1(RNAi)* embryo. Inset, a magnified view of ABplpappap or its descendants. The embryos shown carry the transgenes *stIs10026* and *nIs861*. A, anterior; V, ventral. Scale bar, 10 µm. (E) Ratio of maximum area (see Materials and methods) of ABplpappap to that of its sister in *ced-3(lf)* embryos with RNAi against control, *pig-1* or *gpr-1*. Error bars, standard deviation. ****, p<0.0001 per ordinary one-way ANOVA of log of ratios. n=5 embryos for each RNAi treatment. (F) Quantification of the maximum area (see Materials and methods) of ABplpappap in control embryos, *pig-1(RNAi)* embryos and *grp-1(RNAi)* embryos. Error bars, standard deviation. **, p<0.01; ****, p<0.0001 per ordinary one-way ANOVA. n=5 embryos for each RNAi treatment. (G) Ratio of maximum area (see Materials and methods) of ABplpappap to that of its sister in *ced-3(lf)* embryos with RNAi against control, *cye-1* or *cdk-2*. Error bars, standard deviation. n.s., not significant per ordinary one-way ANOVA of log of ratios. n=4 embryos for each RNAi treatment. (H) Quantification of the maximum area (see Materials and methods) of ABplpappap in control embryos, *cye-1(RNAi)* embryos and *cdk-2(RNAi)* embryos. Error bars, standard deviation. n.s., not significant per ordinary one-way ANOVA. n=4 embryos for each RNAi treatment. (I) Penetrances of the Tex phenotype and number of animals with one, two or three excretory cells produced by RNAi against genes identified from genetic screens for defective extrusion. The *ced-3(lf)* strain used for this experiment carries the transgene *nIs433*.

Next, we investigated why ABplpappap completed the cell cycle in *pig-1(RNAi)* and *grp-1(RNAi)* embryos. In several cell lineages, such as the Q neuroblast cell lineage, the genes *pig-1* and *grp-1* are required for unequal cell divisions that generate apoptotic cells (Cordes *et al*., 2006; Teuliere *et al*., 2014). Consistent with their function in controlling unequal cell division, RNAi against each of the genes *pig-1* and *grp-1* perturbed the ratio of ABplpappap’s size to that of its sister and generated an abnormally large ABplpappap (Figures 5E,F; Supplemental Figures 6A,D,E). These findings indicated that unequal cell division plays an important role in producing the S-phase arrest that precedes cell extrusion. However, despite the requirement for *cye-1* and *cdk-2* to produce this S-phase arrest (Figures 3B, 4B), RNAi against *cye-1* or *cdk-2* did not affect unequal cell division. Neither the size of ABplpappap relative to that of its sister nor the absolute size of ABplpappap showed a difference among *cye-1(RNAi)*, *cdk-2(RNAi)* and control embryos (Figures 5G,H; Supplemental Figure 6A,B,C). Together, our data indicate that genes required for cell extrusion function to produce the S-phase arrest preceding extrusion either by promoting S-phase entry (via *cye-1* and *cdk-2*) or by preventing cell-cycle completion (via *pig-1* and *grp-1*).

Another difference consistent with the distinct ways by which the cell-cycle genes and unequal-cell-division genes promote cell extrusion was observed in the Tex phenotype caused by RNAi against these genes. Some *pig-1(RNAi)* and *grp-1(RNAi)* animals with the Tex phenotype displayed two supernumerary excretory cells (Figure 5I), presumably because both daughters of ABplpappap adopted the excretory-cell in these animals. By contrast, *cye-1(RNAi)* and *cdk-2(RNAi)* animals with the Tex phenotype displayed only one supernumerary excretory cell (Figure 5I), presumably because ABplpappap did not enter and then complete the cell cycle but rather differentiated directly into an excretory cell.

In short, these findings indicate that the unifying feature of all genes required for cell extrusion is that they function to produce the S-phase arrest observed in cells to be extruded, supporting the conclusion that S-phase arrest is a key requirement of cell extrusion. Either preventing cell-cycle entry or bypassing the S-phase arrest to complete cell division prevented cell extrusion in developing *C. elegans* embryos.

### S-phase arrest drives cell extrusion from mammalian epithelial cultures

Since S-phase arrest is the central feature of cell extrusion by *C. elegans* embryo, we asked if S-phase arrest also induces mammalian cell extrusion. We used a monolayer of Madin-Darby Canine Kidney (MDCK) cells as a model of mammalian epithelia and hydroxyurea (HU) as a chemical agent for inducing S-phase arrest. MDCK cells are a simple epithelial system for studies of mammalian cell extrusion in culture (reviewed by Ohsawa *et al*., 2018). HU causes S-phase arrest by inhibiting the enzyme ribonucleotide reductase and depleting deoxyribonucleotides during DNA replication, resulting in stalled DNA replication forks (Timson, 1975; Bianchi *et al*., 1983). We treated MDCK monolayers with either 2 mM HU or vehicle (negative control) for up to 24 h and obtained time-lapse micrographs of the monolayers from this period to assess cell extrusion. Strikingly, cells extruded apically from the MDCK monolayer into the culture medium were three to four times higher in number after HU treatment when compared to the number of cells extruded apically after vehicle treatment (Figure 6A, 6B; Movie 6,7). Next we used MDCK-Fucci cells to determine the cell cycle phase distribution of HU- and vehicle-treated extruded cells (Streichan *et al*., 2014). MDCK-Fucci cells produce a fluorescent signal that varies with the phase of the cell cycle (G0/G1-Red, S/G2/M-Green; Sakaue-Sawano et al., 2008). As expected, most of the HU-treated extruded cells displayed a green fluorescent signal (Figure 6C), indicating that they were in a cell cycle phase subsequent to the onset of DNA replication. We noted that stochastically extruded cells (from vehicle treatment) mostly exhibited red fluorescence indicative of the G0 or G1 phase (Figure 6D), consistent with the phase of cells that are naturally extruded from post-mitotic zones, such as the tips of intestinal villi (Carroll *et al*., 2018; Eisenhoffer *et al*., 2012).

**Figure 6.**
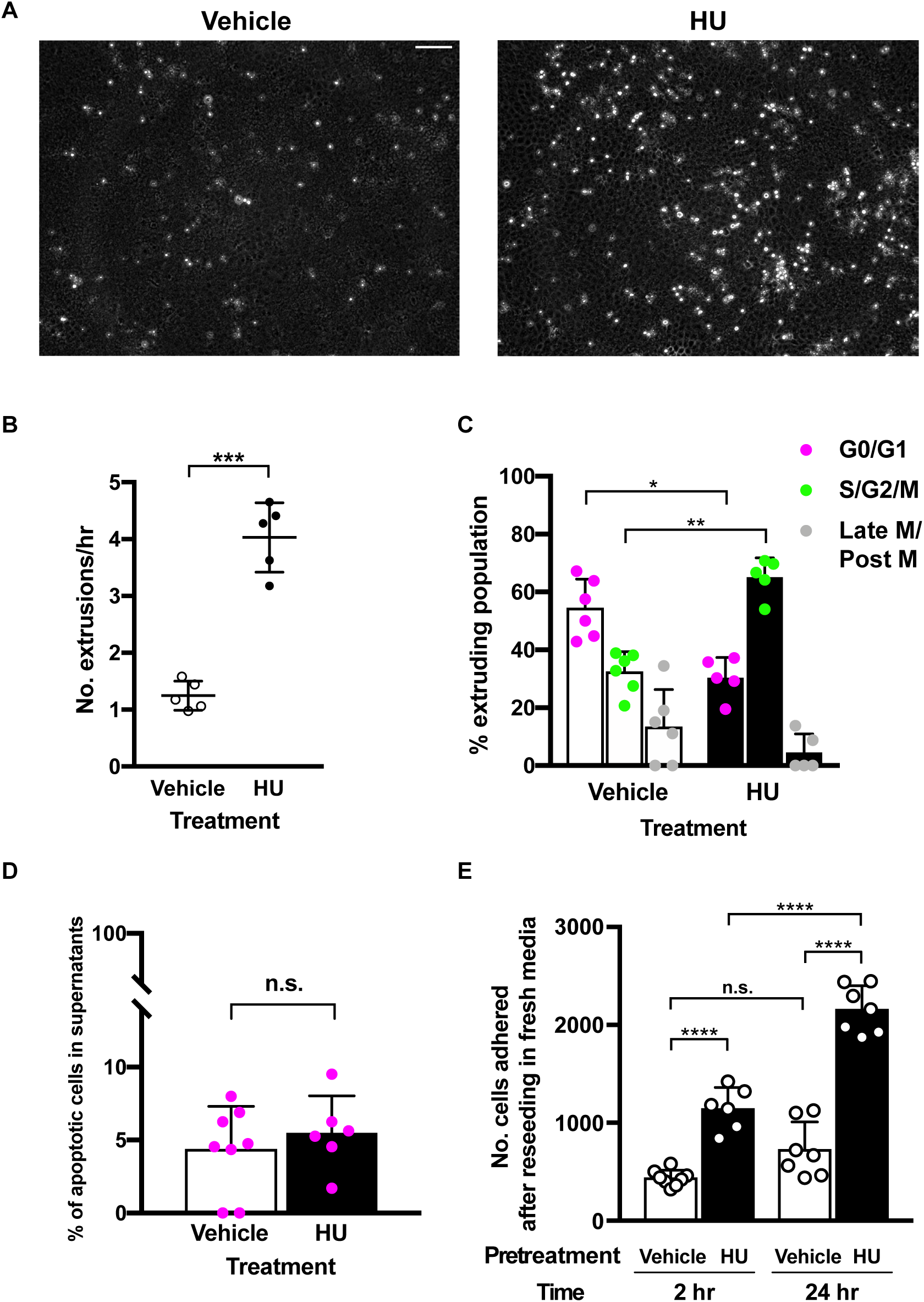
S-phase arrest is sufficient for cell extrusion from a simple mammalian epithelial layer. (A) Representative images from time-lapse videos of mammalian MDCK cell monolayers exposed to either vehicle or 2 mM hydroxyurea (HU) at 22 h of exposure. Extruded cells can be identified as bright, white, rounded spots rising from the epithelial plane. Many more extruded cells are observed after HU treatment as compared to vehicle treatment. Scale bar, 100 µm. (B) Graph showing average number of cell extrusions per h in vehicle-treated and 2 mM HU-treated MDCK monolayers. Each data point represents a separate experiment conducted for up to 24 h normalized by duration of experiment for comparison. Error bars, standard deviation. ***, p < 0.001 per Welch’s two-tailed t-test. n=5 measurements experimental replicates for each condition. (C) Graph showing cell cycle phase distribution of extruded MDCK-FUCCI cells after HU treatment or vehicle treatment. Each data point represents a separate experiment. Error bars, standard deviation. *, p<0.05; **, p<0.01 per Kruskal-Wallis test. (D) Graph showing percentage of cells extruded after vehicle treatment or HU treatment that are apoptotic as per Trypan blue uptake. Each data point represents a separate experiment. Error bars, standard deviation. n.s., not significant per Mann-Whitney test. (E) Graph showing the number of HU-treated or vehicle-treated extruded cells that adhered at 2 h and 24 h after reseeding in fresh media. Each data point represents a separate experiment. Error bars, standard deviation. n.s., not significant; ****, p<0.0001 per standard one-way ANOVA.

Since HU is known to increase the rate of apoptosis (Timson, 1975), and agents that promote apoptosis increase the rate of cell extrusion (Andrade and Rosenblatt, 2011), we examined the role of apoptosis in HU-induced cell extrusion. Surprisingly, the fraction of extruded MDCK cells that were apoptotic was not increased by HU treatment (Figure 6D), indicating that HU-induced cell extrusion is not a consequence of an increase in the rate of apoptosis. In addition, we reseeded the extruded MDCK cells in fresh media to measure their viability following HU treatment. We found that the number of viable adherent cells at 2 hours post reseeding was proportional to the number of extrusions for both the HU- and vehicle-treated groups (Figure 6E). Additionally, the number of HU-treated cells doubled at 24 hours post reseeding when compared to 2 hours post reseeding (Figure 6E). Thus, cells extruded by HU treatment were not only viable but also able to resume and complete the cell cycle.

Taken together, the above findings indicate that S-phase arrest drives the extrusion of cells from mammalian epithelia and establish that cell extrusion caused by S-phase arrest is an evolutionarily conserved phenomenon.

## DISCUSSION

Here we report that cell cycle S-phase arrest is a cell-intrinsic trigger of cell extrusion and can induce the extrusion of cells of organisms from two divergent branches of the phylogenetic tree - nematodes and mammals. Using RNAi screens and transgenic cell-cycle reporters, we determined in developing *C. elegans* embryos that perturbations that prevented S-phase arrest blocked cell extrusion. As summarized in Figure 7A, cells destined for extrusion from *ced-3(lf)* embryos are always the smaller daughters of unequal cell divisions. These smaller daughter cells enter S phase of the cell cycle and undergo S-phase arrest, likely because of a deficiency in the energetic and metabolic resources required for DNA synthesis (e.g., nucleotides, replication proteins, etc.). Thus, either bypassing S-phase arrest (by perturbing the process of unequal cell division) and hence completing cell division or preventing S-phase arrest (by blocking cell-cycle entry) prevents cell extrusion in *C. elegans* embryos.

**Figure 7.**
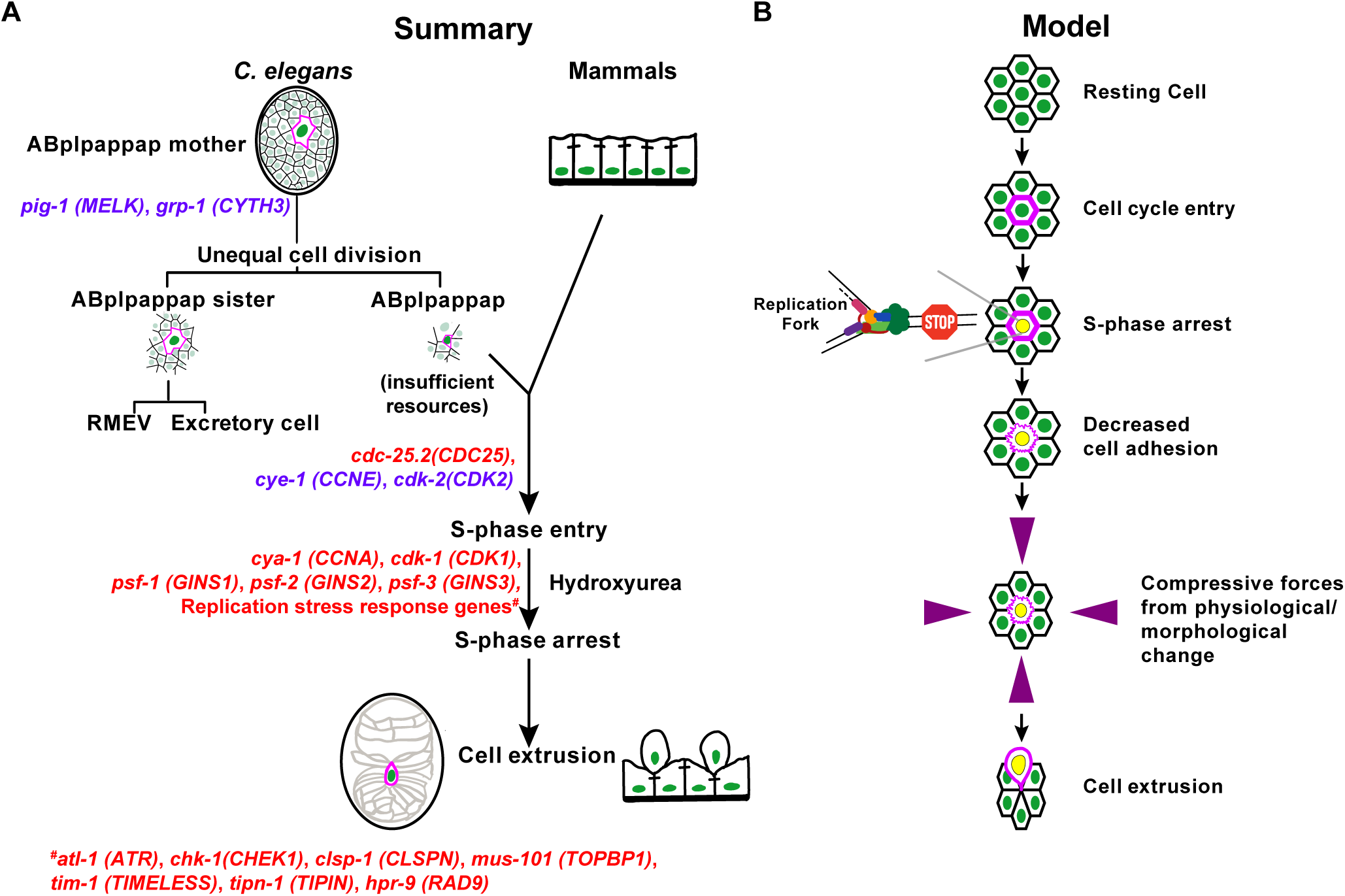
Summary and Model: S-phase arrest drives cell extrusion. (A) A summary of the genes required for cell extrusion by *C. elegans*, their mammalian homologs (in parentheses) and their associated biological processes that precede the S-phase arrest that drives the cell extrusion of ABplpappap (and other extruded cells) from *C. elegans* embryos. Treatment of mammalian cells (MDCK) with HU chemically produces an S-phase arrest in cells that drives their extrusion from simple epithelial layers, as shown. At each step preceding cell extrusion by *C. elegans*, genes with function we demonstrated to occur at the corresponding step are shown in purple, and genes with probable function at that step are shown in red. Relevant cells in the *C. elegans* embryos are outlined in magenta. The horizontal hyphen-like lines connecting mammalian cells indicate adhesion junctions. (B) Model: A cell that enters the cell cycle (marked by magenta cell boundary) but arrests in S phase (marked by yellow nucleus) will have lowered cell adhesion (marked by wavy magenta cell boundary). When a cell with lowered cell adhesion caused by S-phase arrest experiences morphological or physiological forces (marked by purple arrows) that cause a squeezing-like effect, the cell gets extruded.

To test the generality of our findings, we used mammalian epithelia treated with HU and showed that S-phase arrest is sufficient to promote cell extrusion. Thus, cell extrusion triggered by S-phase arrest is an evolutionarily conserved mechanism of cell elimination. These observations also demonstrate that cells can be extruded from mitotically active mammalian epithelial tissues. Previous studies of mammalian cell extrusion focused on extrusion from post-mitotic tissues (Rosenblatt *et al*., 2001; Eisenhoffer *et al*., 2012; Gudipaty *et al*., 2017; Saw *et al*., 2017; Kocgozlu *et al*., 2016) or oncogene-driven extrusion from growth-suppressing epithelial environments (Anton *et al*., 2018; Hogan *et al*., 2009; Kajita *et al*., 2010; Leung and Brugge, 2012; Slattum *et al*., 2014; Wu *et al*., 2014).

Our mechanistic model for the evolutionarily conserved process by which S-phase arrest promotes cell extrusion is presented in Figure 7B. In this model, a mitotically active cell destined for extrusion (i) enters the cell cycle, (ii) arrests in S phase, (iii) loses cell adhesion (see below), and (iv) is extruded as a result of reduced adhesion and forces generated by external morphological or physiological processes.

### Why are S-phase arrested cells susceptible to extrusion?

We observed previously that cells extruded by *C. elegans* embryos do not express the classical E-cadherin HMR-1 and other cell-adhesion molecules (Denning *et al*., 2012). The absence of such adhesion molecules likely allows cells to be squeezed out of the embryo by morphological forces generated by migrating hypodermal cells and neighboring neuroblasts during ventral enclosure (Chisholm and Hardin, 2005; Wernike *et al*., 2016). Consistent with this view, the loss of E-cadherin-mediated adhesion caused by cleavage of the extracellular part of a cell’s E-cadherin molecules is sufficient to drive cell extrusion from an MDCK monolayer (Grieve and Rabouille, 2014). We speculate that a signaling pathway initiated by S-phase arrest downregulates the expression of cell adhesion molecules in cells destined for extrusion. Indeed, in HeLa cell cultures, cell adhesion increases during S phase via the activity of the cell-cycle regulator CDK1 and decreases later in the cell cycle upon inhibition of CDK1 by Wee1 (Jones *et al*., 2018). Interestingly, activation of the replication stress response, which is required for cell extrusion in *C. elegans* (Figure 1C, 4L), blocks CDK1 activity (Jin *et al*., 2003; Mailand *et al*., 2002; Xiao *et al*., 2003). Furthermore, HU-mediated S-phase arrest also inactivates CDK1 in MDCK cells (Anton *et al*., 2018). We therefore propose that a reduction in CDK1 activity following S-phase arrest decreases cell adhesion, thereby facilitating the extrusion of cells subjected to external forces from cellular neighbors.

### Extrusion of S-phase arrested cells is likely tumor-suppressive

Replication forks under prolonged S-phase arrest can collapse and produce DNA damage, genomic rearrangements and ploidy defects, all of which are associated with oncogenesis (reviewed by Gaillard *et al*., 2015). The human genes that promote replication stress and S-phase arrest are frequently amplified, overexpressed or activated by mutations in tumors (Otto and Sicinski, 2017). Cells in such tumors experience persistent replication stress that can lead to S-phase arrest (Gaillard *et al*., 2015). Hence, tumor cells and cells with oncogenic potential might well be eliminated via cell extrusion, in which case the extrusion of cells arrested in S phase would be tumor-suppressive. We propose that cell extrusion driven by S-phase arrest is a checkpoint mechanism that functions to eliminate cells at all stages of the oncogenic transformation process, ranging from precancerous cells in S-phase arrest to tumor cells in a malignant tumor.

### Subversion of cell extrusion driven by S-phase arrest might contribute to metastasis

Inactivation of either S1P_2_ or the tumor suppressor APC or expression of oncogenic K-Ras changes the direction of cell extrusion from apical, which favors cell elimination by extrusion into the lumen, to basal, which favors dissemination of extruded cells to surrounding tissue (Gu *et al*., 2015; Marshall *et al*., 2011; Slattum *et al*., 2014). Mutations in these genes are hallmarks of metastatic tumors. While cell extrusion caused by S-phase arrest likely can suppress tumor development, the same mechanism of cell extrusion might paradoxically promote cancer metastasis if the extrusion direction changes from apical to basal. We observed that cells subjected to HU-mediated extrusion failed to die (Figure 6D) and instead were capable of reentering the cell cycle and proliferating (Figure 6E). Thus, basal extrusion of cells arrested in S-phase might facilitate metastasis by disseminating live tumor cells arrested in S phase to other tissues and organs. We propose that mutations in genes, such as S1P_2_, APC and K-Ras, promote metastasis by facilitating the basal extrusion and spread of tumor cells arrested in S phase.

In summary, we have discovered a novel conserved mechanism that links a cell-cycle vulnerability to the process of cell extrusion. We suggest that cell extrusion mediated by S-phase arrest is a mechanism of cell elimination common to all metazoa. These findings have implications for the field of cancer biology, as cell extrusion caused by S-phase arrest likely regulates both the survival and spread of tumor cells.

## ACKNOWLEDGMENTS

We thank S. van den Heuvel for providing strains; the CGC, which is funded by NIH Office of Research Infrastructure Programs (P40 OD010440), for providing some strains; N. An and T. Ljungars for strain management; S. Luo, S.R. Sando, A. Doi, and A. Corrionero and other members of the Horvitz laboratory for helpful discussions, and D. Ghosh, C.L. Pender, J.N. Kong, M.G. Vander Heiden, P.W. Reddien, R.O. Hynes for suggestions regarding the manuscript. V.K.D. was a Howard Hughes Medical Institute International Student Research fellow. C.P.P. is the recipient of a Human Frontiers Science Program postdoctoral fellowship (LT000654/2019-L). J.R. and C.P.P. were funded by King’s College London startup funds. H.R.H. is the David H. Koch Professor of Biology at MIT and an Investigator at the Howard Hughes Medical Institute.

## AUTHOR CONTRIBUTIONS

H.R.H. supervised the project. V.K.D. and H.R.H. conceptualized the project. V.K.D. and H.R.H. designed the experiments that used *C. elegans*. V.K.D. and R.D. performed the experiments that used *C. elegans*. V.K.D., R.D. and D.P.D. generated reagents. C.P. and J.R. designed the experiments that used mammalian cells. C.P. performed the experiments that used mammalian cells. V.K.D., D.P.D. and H.R.H. wrote the original manuscript draft. All authors contributed to data analysis, interpretation, and reviewing and editing of the manuscript.

---

**Supplemental Table 1.** Penetrance of the Tex phenotype produced by RNAi against cell cycle genes (and non-cell-cycle cyclins and CDKs) in *ced-3(lf)* animals. Tex penetrance produced by each of the RNAi clones in the cell-cycle RNAi library, the number of animals counted for each RNAi clone and whether or not the RNAi clone produced extensive lethality are shown. Genes corresponding to RNAi clones that produced more than 9% penetrance of the Tex phenotype are in bold. *cyb-3* did not fit this criterion, as extensive lethality prevented the counting of sufficient number of animals to assign significance. Some cyclins and CDKs that function outside the cell cycle were included in this library and served as negative controls.

| RNAi target | Mammalian Homolog | %Tex | n | extensive lethality? | RNAi target | Mammalian Homolog | %Tex | n | extensive lethality? |
| --- | --- | --- | --- | --- | --- | --- | --- | --- | --- |
| empty vector | - | 1 | 159 | N | *cic-1* | CCNC | 1 | 158 | N |
| ***atl-1*** | ATR | 10 | 509 | N | *cyd-1* | CCND | 0 | 111 | N |
| *cdc-14* | CDC14 | 1 | 168 | N | ***cye-1*** | CCNE | 89 | 133 | Y |
| *cdc-25.1* | CDC25 | 0 | 111 | Y | *cyh-1* | CCNH | 0 | 119 | Y |
| ***cdc-25.2*** | CDC25 | 12 | 159 | N | *cyl-1* | CCNL | 1 | 106 | Y |
| *cdc-25.3* | CDC25 | 2 | 168 | N | *cyy-1* | CCNY | 1 | 108 | N |
| *cdc-25.4* | CDC25 | 0 | 183 | N | *dpl-1* | TFDP1 | 0 | 167 | N |
| ***cdk-1*** | CDK1 | 15 | 61 | Y | *efl-1* | E2F | 0 | 141 | N |
| *cdk-11.1* | CDK11 | 0 | 175 | N | *emb-27* | CDC16 | 0 | 107 | Y |
| *cdk-11.2* | CDK11 | 1 | 132 | N | *emb-30* | ANAPC4 | 0 | 130 | Y |
| *cdk-12* | CDK12 | 0 | 167 | N | *fzr-1* | FZR1 | 1 | 146 | N |
| ***cdk-2*** | CDK2 | 65 | 181 | N | *fzy-1* | CDC20 | 0 | 130 | Y |
| *cdk-4* | CDK4 | 1 | 193 | N | *hpr-17* | RAD17 | 5 | 214 | N |
| *cdk-5* | CDK5 | 0 | 155 | N | *hus-1* | HUS1 | 0 | 132 | N |
| *cdk-7* | CDK7 | 0 | 130 | Y | *lin-15* | - | 1 | 104 | N |
| *cdk-8* | CDK8 | 1 | 186 | N | ***lin-23*** | βTrCP | 23 | 147 | N |
| *cdk-9* | CDK9 | 1 | 155 | Y | *lin-35* | Rb | 0 | 134 | N |
| *cdt-1* | CDT1 | 1 | 147 | Y | *lin-36* | - | 0 | 111 | N |
| ***chk-1*** | CHEK1 | 10 | 164 | Y | *lin-9* | LIN9 | 1 | 125 | N |
| *cit-1.1* | CCNT1/2 | 0 | 105 | N | *mat-1* | CDC27 | 0 | 80 | Y |
| *cit-1.2* | CCNT1/2 | 1 | 244 | N | *mat-2* | ANAPC1 | 1 | 163 | Y |
| *cki-1* | CDKN1 | 0 | 125 | N | *mat-3* | CDC23 | 0 | 108 | Y |
| *cki-2* | CDKN1 | 1 | 216 | N | *mdf-1* | MAD1L1 | 3 | 112 | N |
| *clk-2* | TELO2 | 1 | 156 | N | *mdf-2* | MAD2L1 | 1 | 111 | N |
| *cul-1* | CUL1 | 5 | 81 | Y | *mrt-2* | RAD1 | 0 | 114 | N |
| *cul-2* | CUL2 | 0 | 21 | Y | *rnr-1* | RRM1 | 5 | 103 | Y |
| *cul-3* | CUL3 | 5 | 151 | Y | *san-1* | BUB1B | 1 | 145 | N |
| *cul-4* | CUL4 | 0 | 141 | N | *wee-1.1* | PKMYT1 | 1 | 159 | N |
| ***cya-1*** | CCNA | 20 | 309 | N | *wee-1.3* | PKMYT1 | 0 | 60 | Y |
| *cyb-1* | CCNB1 | 4 | 136 | N |  |  |  |  |  |
| *cyb-2.1* | CCNB2 | 1 | 102 | N |  |  |  |  |  |
| *cyb-2.2* | CCNB2 | 0 | 144 | N |  |  |  |  |  |
| *cyb-3* | CCNB3 | 14 | 7 | Y |  |  |  |  |  |

**Supplemental Table 2.**
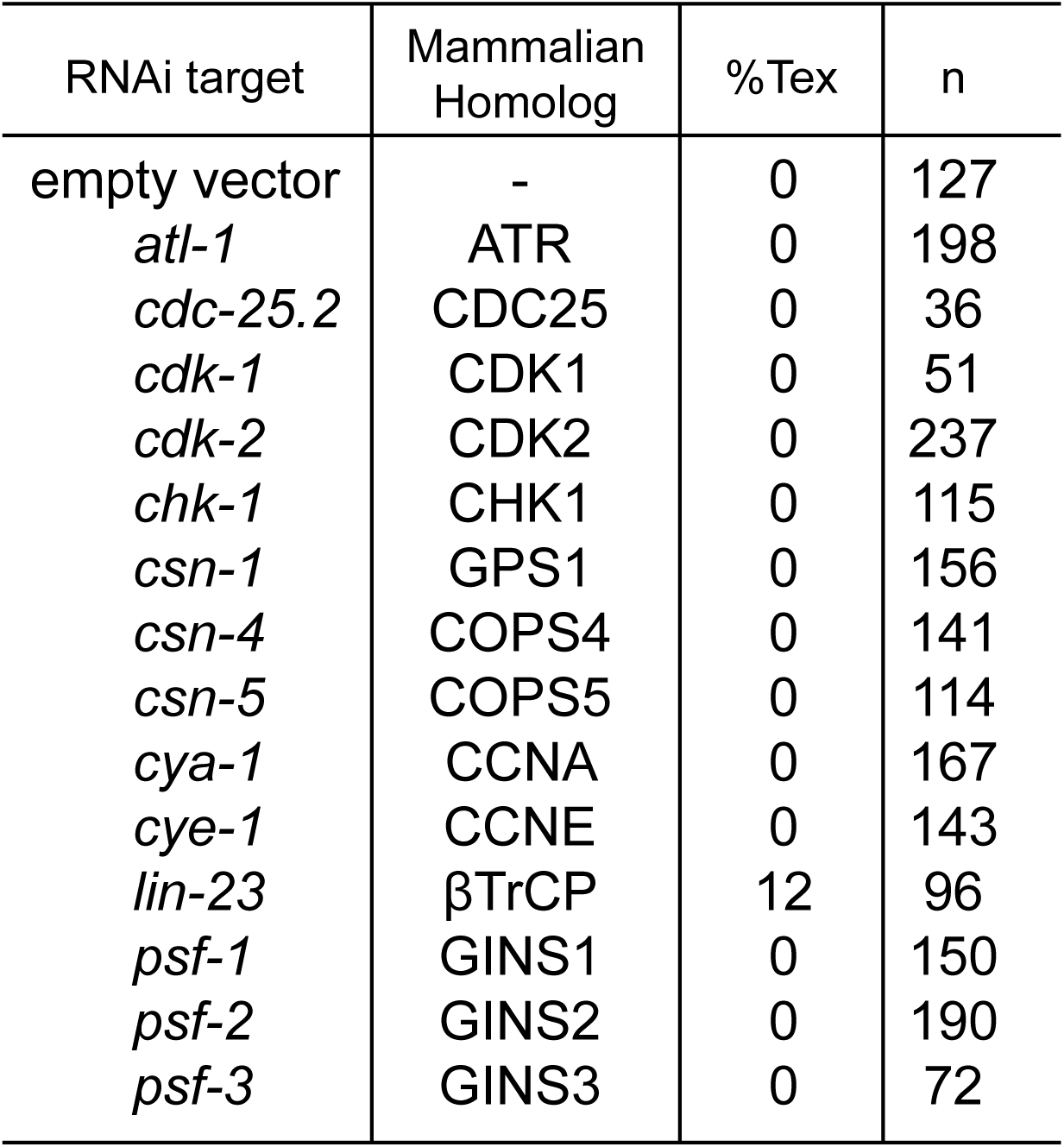
Penetrance of the Tex phenotype produced in wild-type animals by RNAi against cell cycle genes with potential roles in cell extrusion. The Tex penetrance produced in wild-type animals by RNAi clones against cell cycle genes that might be involved in cell extrusion (based on the corresponding Tex penetrance in *ced-3(lf)* animals) is provided. *Bona fide* candidates for cell extrusion regulation should not produce a Tex phenotype in wild-type animals, as cell extrusion does not occur in wild-type worms. A Tex phenotype in wild-type animals could occur from other effects of RNAi against cell cycle genes, such as excessive proliferation leading to multiple excretory cells. Such proliferation is likely the case for *lin-23*, as RNAi against *lin-23* has been previously described to cause excessive proliferation (Kipreos *et al*., 2000). The 13 other genes are good candidates to be regulators of cell extrusion by the criterion of dependence of the Tex phenotype on the loss of function of *ced-3*.

**Supplemental Figure 1.**
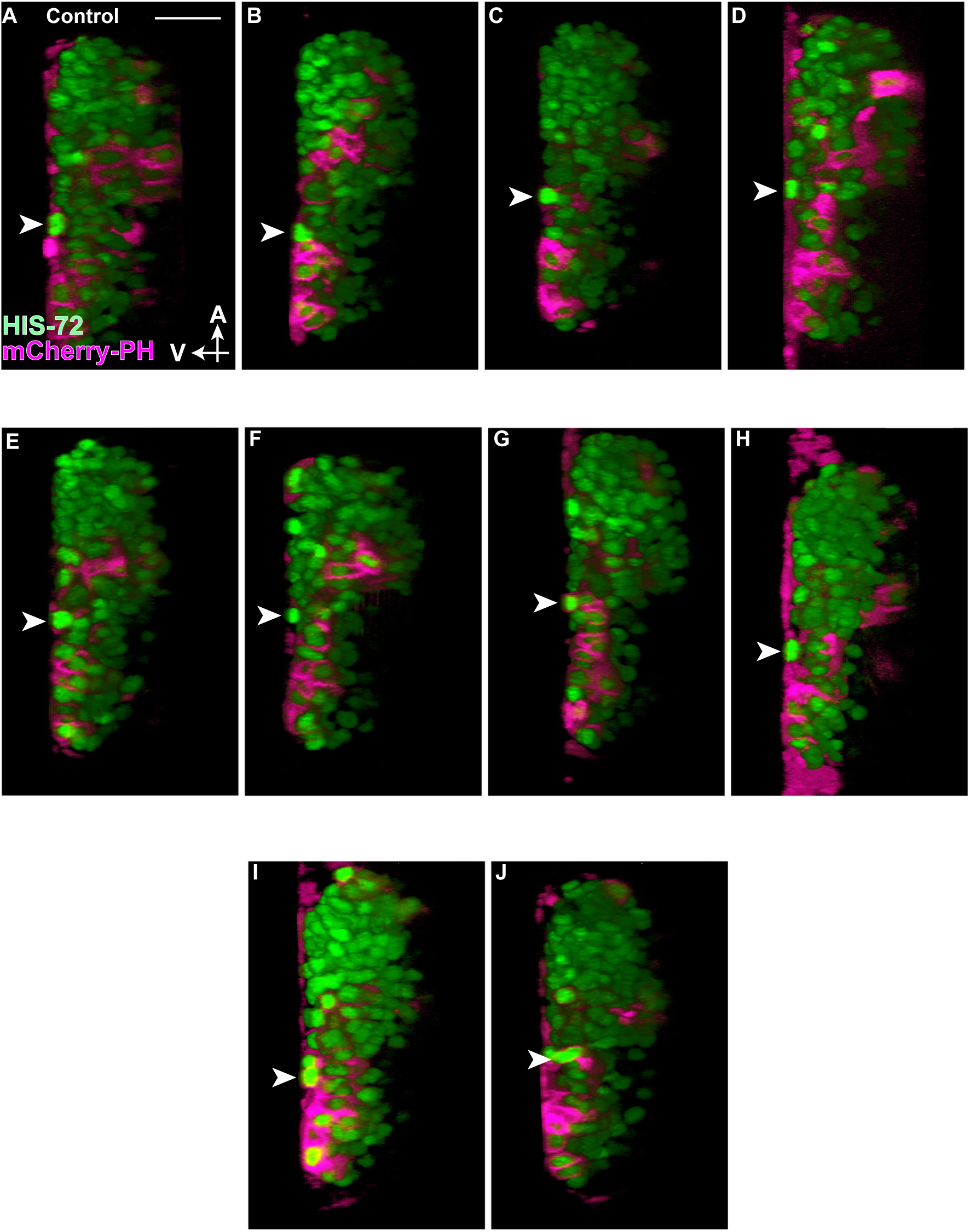
ABplpappap is extruded by control embryos. (A-J) Virtual lateral sections of embryos through the ABplpappap cell (arrowhead) in (A-J) 10 control embryos show (A-I) ABplpappap is extruded in 9 of 10 embryos and (J) is not extruded in 1 of 10 embryos. The embryos shown carry the transgenes *stIs10026* and *nIs861*. A, anterior; V, ventral. Scale bar, 10 µm.

**Supplemental Figure 2.**
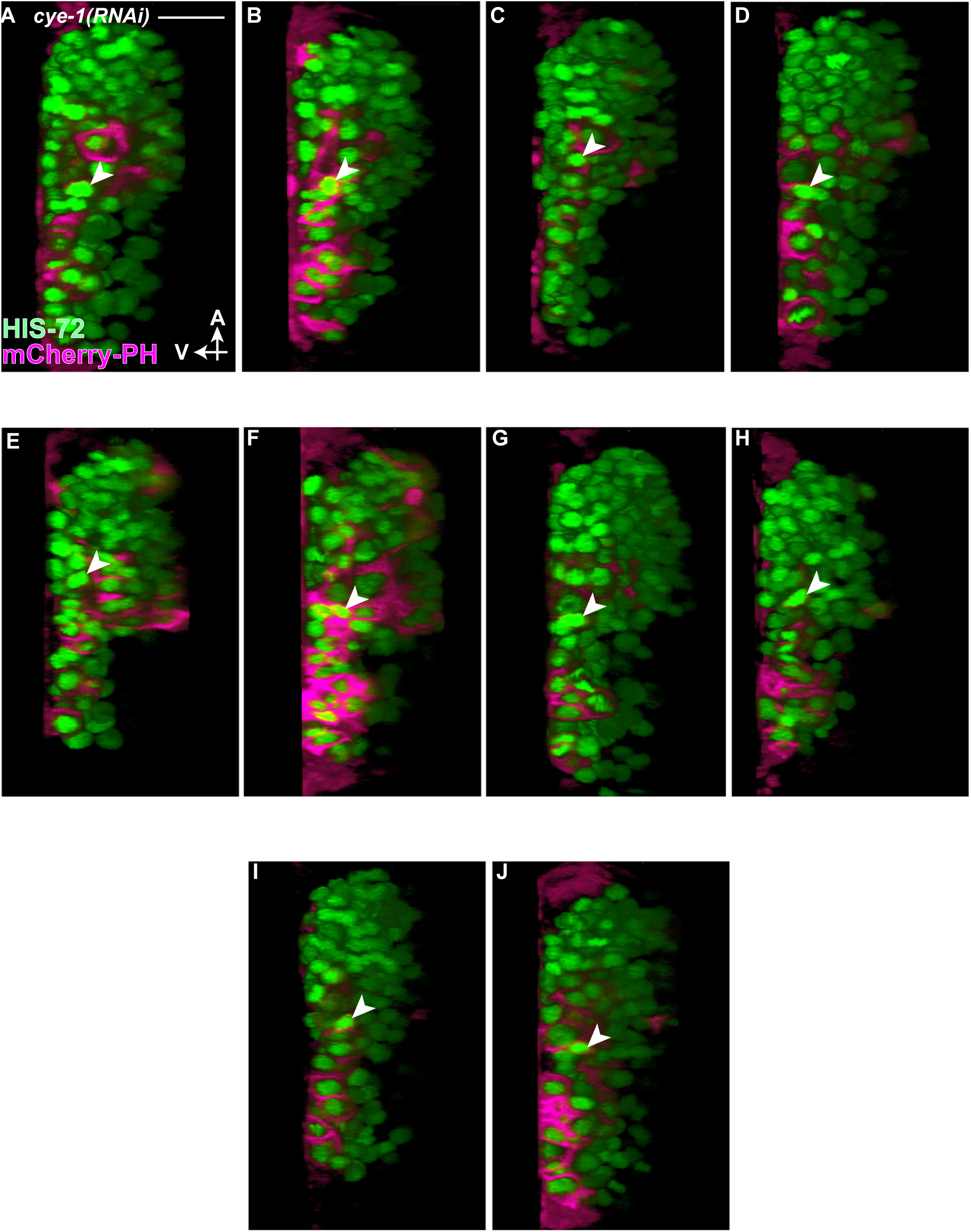
ABplpappap is not extruded by *cye-1(RNAi)* embryos. (A-J) Virtual lateral sections of embryos through the ABplpappap cell (arrowhead) in (A-J) 10 *cye-1(RNAi)* embryos show that ABplpappap is not extruded in 10 of 10 embryos. The embryos shown carry the transgenes *stIs10026* and *nIs861*. A, anterior; V, ventral. Scale bar, 10 µm.

**Supplemental Figure 3.**
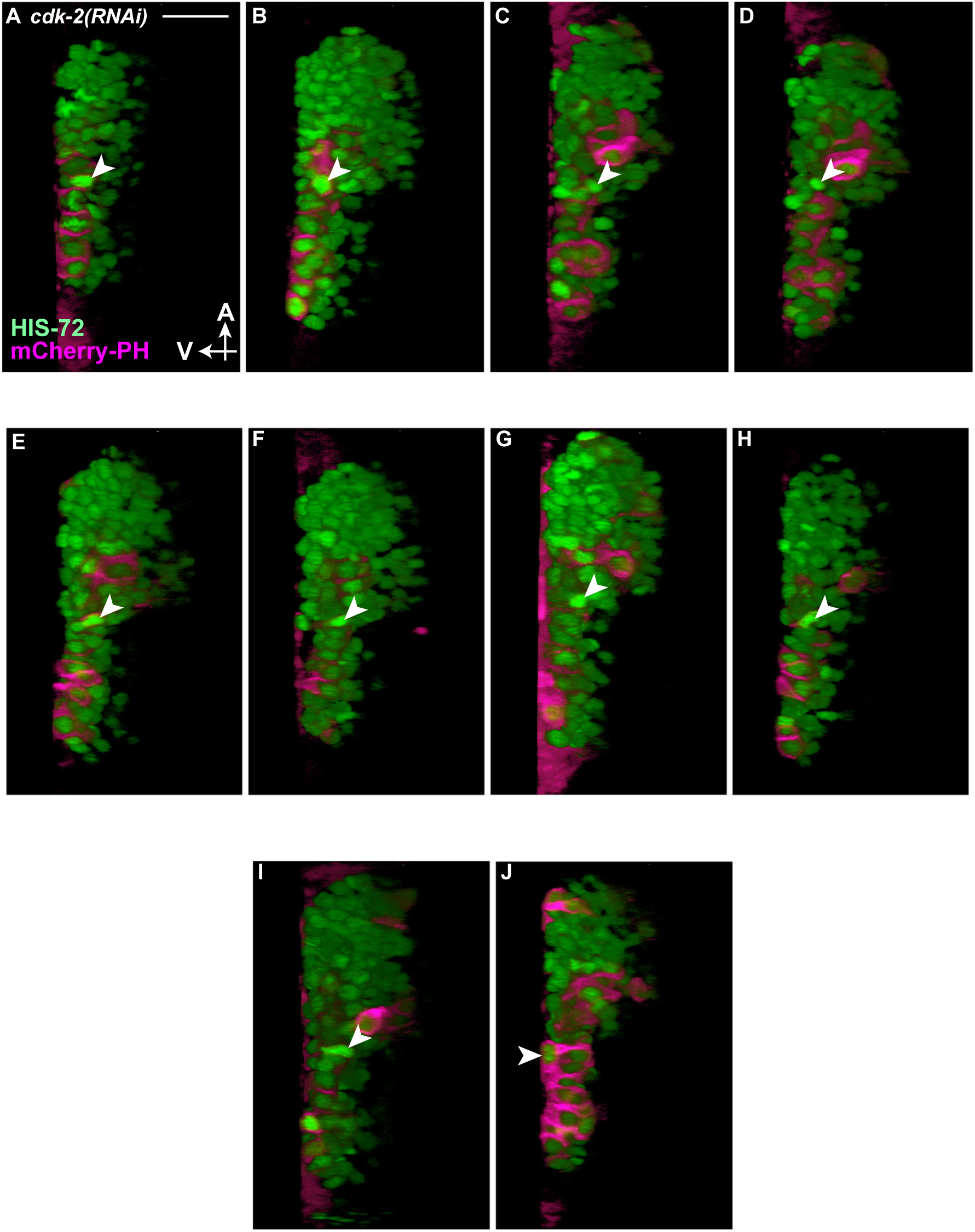
ABplpappap is not extruded by *cdk-2(RNAi)* embryos. (A-J) Virtual lateral sections of embryos through the ABplpappap cell (arrowhead) in (A-J) 10 *cdk-2(RNAi)* embryos show that (A-I) ABplpappap is not extruded in 9 of 10 embryos and (J) is extruded in 1 of 10 embryos. The embryos shown carry the transgenes *stIs10026* and *nIs861*. A, anterior; V, ventral. Scale bar, 10 µm.

**Supplemental Figure 4.**
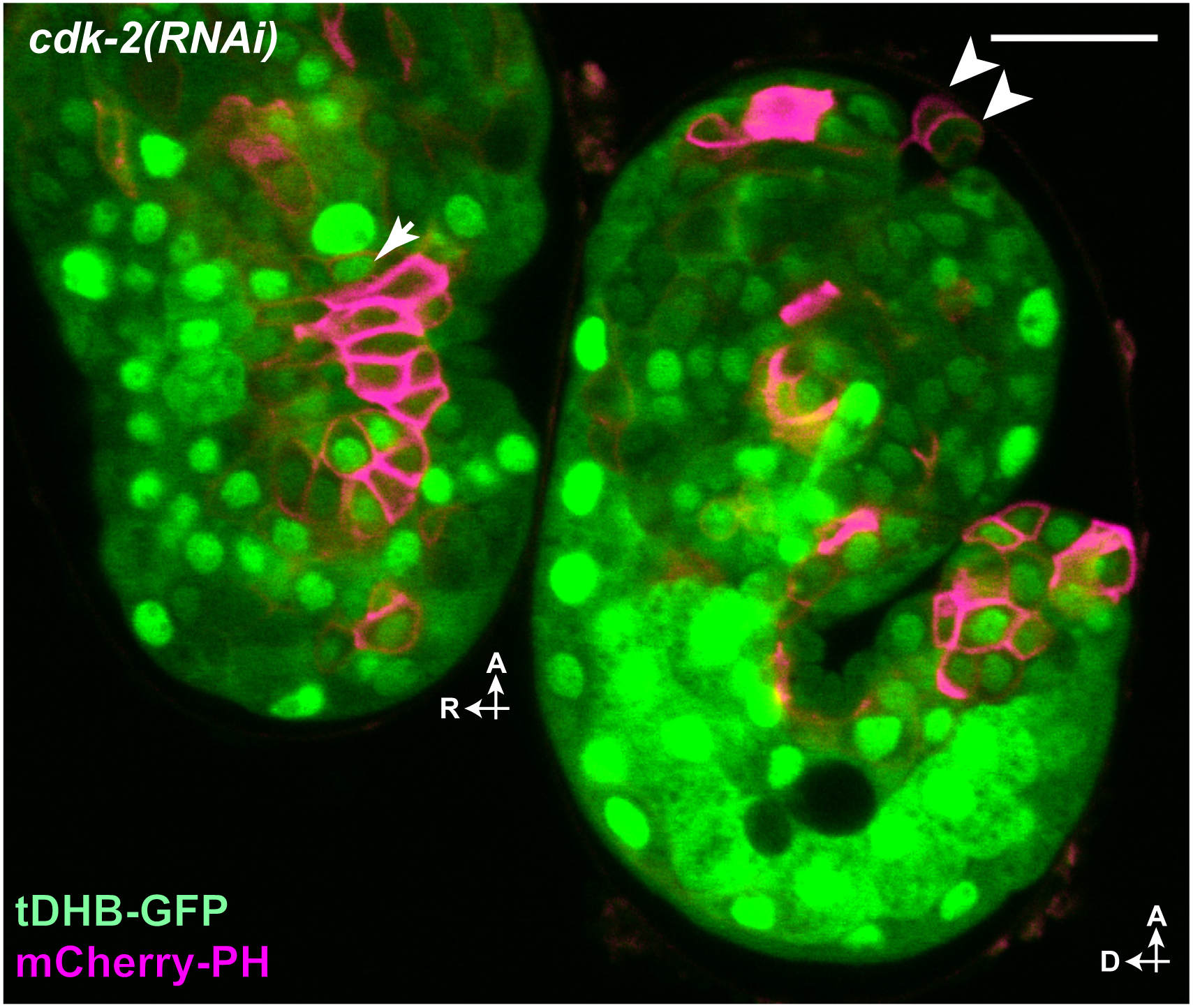
Cells extruded by *cdk-2(RNAi)* embryos enter the cell cycle. Micrograph of two *cdk-2(RNAi)* embryos expressing tDHB-GFP shows a nuclear-enriched localization of tDHB-GFP in an ABplpappap cell (arrow) that was not extruded by the first embryo but presumably nuclear-depleted tDHB-GFP in two cells that were extruded from the second embryo (arrowheads). The embryos shown carry the transgenes *heSi192* and *nIs861*. A, anterior; D, dorsal; R, right. Scale bar, 10 µm.

**Supplemental Figure 5.**
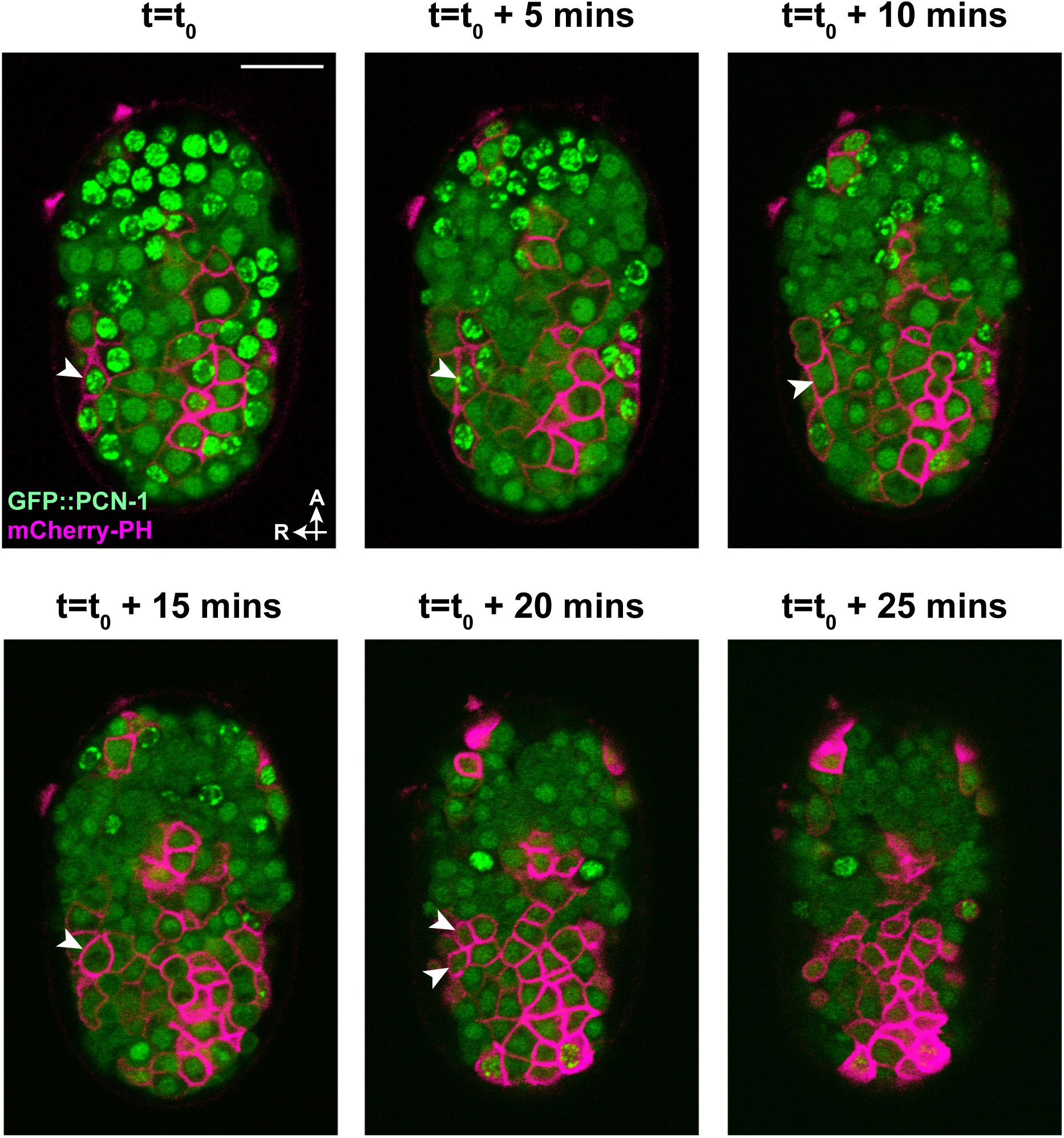
GFP::PCN-1 exhibits a localization pattern coordinated with the cell cycle in embryonic cells on the ventral surface. Time-lapse confocal micrographs of a *ced-3(lf)* embryo obtained at 5-min intervals show the dynamics of GFP::PCN-1 localization in multiple cells on the ventral surface of the embryo. Arrowheads mark a cell that is shown undergoing a complete cell cycle. The embryo shown carries the transgenes *isIs17* and *nIs861*. A, anterior; R, right. Scale bar, 10 µm.

**Supplemental Figure 6.**
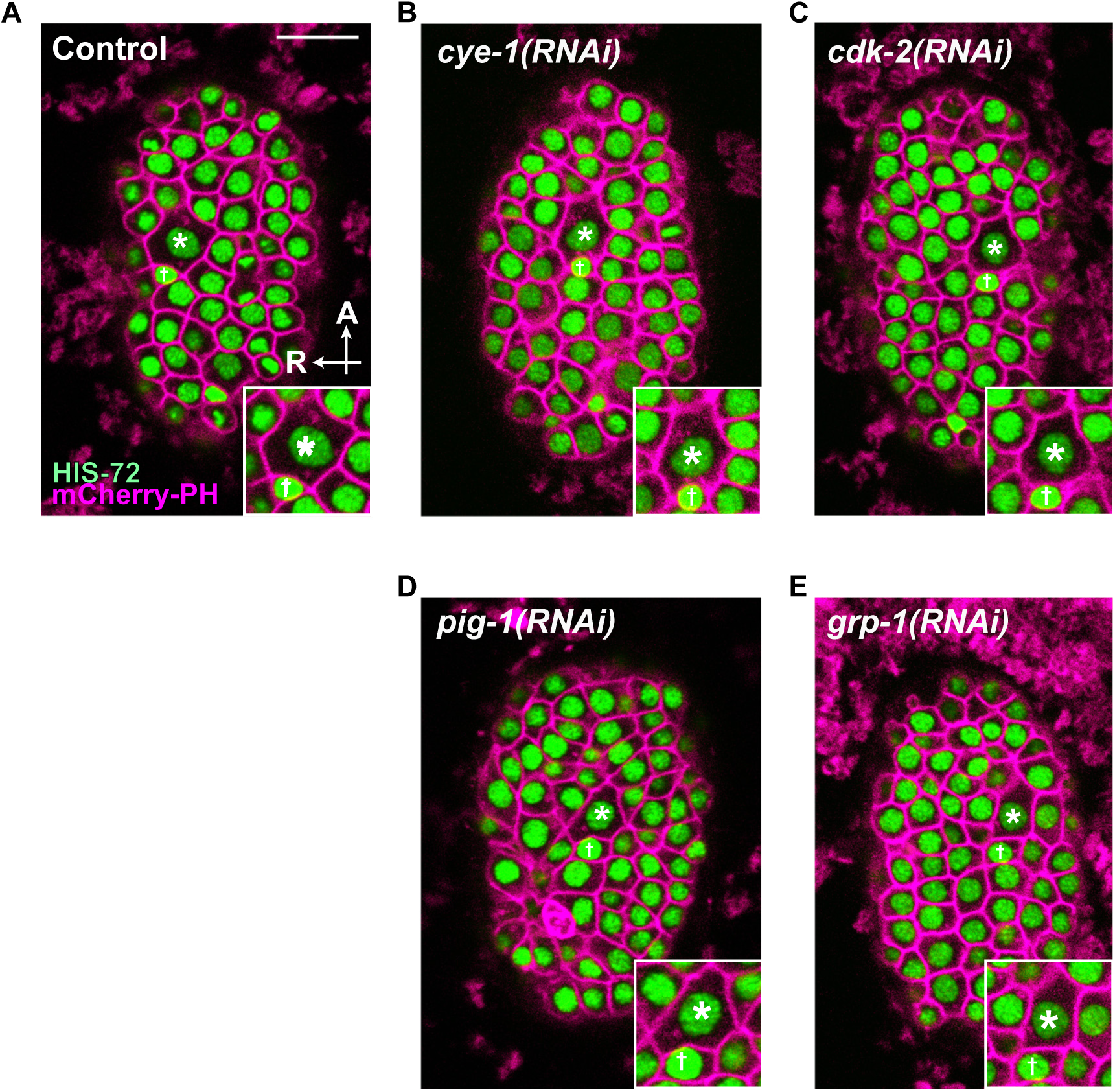
ABplpappa undergoes unequal cell division controlled by *pig-1* and *grp-1* but independent of *cye-1* and *cdk-2*. Micrographs of *ced-3(lf)* embryos expressing nuclear GFP and membrane-localized mCherry in all cells obtained using confocal microscopy show the relative sizes of ABplpappap (†) and its sister, ABplpappaa (*) in a (A) control embryo, (B) *cye-1(RNAi)* embryo, (C) *cdk-2(RNAi)* embryo, (D) *pig-1(RNAi)* embryo and (E) *grp-1(RNAi)* embryo. Inset, a magnified view of ABplpappap(†) and its sister, ABplpappaa(*). All embryos shown carried the transgenes *stIs10026* and *ltIs44[P_pie-1_::mCherry::PH]*, which expresses membrane-localized mCherry in all cells. A, anterior; R, right. Scale bar, 10 µm.

## Supplemental Movies

**Movie 1 Control embryos extrude ABplpappap as it undergoes ventral enclosure**

Time-lapse video of a *ced-3(lf); control(RNAi)* embryo undergoing ventral enclosure over a period of 50 minutes shows ABplpappap (circled at the beginning and end of video) was extruded from this embryo. All nuclei are labeled with GFP and membranes of *egl-1*–expressing cells are labeled with mCherry (magenta). Time-lapse images used for this video were obtained using confocal microcopy. Video playback is at 600x real speed. The embryo shown carried the transgenes *stIs10026* and *nIs632*.

**Movie 2 *cye-1(RNAi)* embryos do not extrude ABplpappap as it undergoes ventral enclosure**

Time-lapse video of a *ced-3(lf); cye-1(RNAi)* embryo undergoing ventral enclosure over a period of 50 minutes shows ABplpappap (circled at the beginning and end of video) was not extruded from this embryo. All nuclei are labeled with GFP and membranes of *egl-1*–expressing cells are labeled with mCherry (magenta). Time-lapse images used for this video were obtained using confocal microcopy. Video playback is at 600x real speed. The embryo shown carried the transgenes *stIs10026* and *nIs632*.

**Movie 3 *cdk-2(RNAi)* embryos do not extrude ABplpappap as it undergoes ventral enclosure**

Time-lapse video of a *ced-3(lf); cdk-2 (RNAi)* embryo undergoing ventral enclosure over a period of 50 minutes shows ABplpappap (circled at the beginning and end of video) was not extruded from this embryo. All nuclei are labeled with GFP and membranes of *egl-1*–expressing cells are labeled with mCherry (magenta). Time-lapse images used for this video were obtained using confocal microcopy. Video playback is at 600x real speed. The embryo shown carried the transgenes *stIs10026* and *nIs632*.

**Movie 4 GFP::PCN-1 shows continuous change in fluorescence intensity during embryonic cell cycles**

Time-lapse video of a *ced-3(lf)* embryo expressing GFP::PCN-1 in all cells shows continuous change in GFP::PCN-1 fluorescence intensity as cells progress through the cell cycle, similar to that observed for early embryonic cell cycles (Brauchle *et al*., 2003). Membranes of *egl-1*–expressing cells are labeled with mCherry (magenta). Time-lapse images used for this video were obtained using confocal microcopy. Video playback is at 180x real speed. The embryo shown carried the transgenes *isIs17* and *nIs861*.

**Movie 5 ABplpappap divides before ventral enclosure and is not extruded in *pig-1(RNAi)* embryos**

Time-lapse video of a *ced-3(lf); pig-1 (RNAi)* embryo undergoing ventral enclosure over a period of 57 minutes shows ABplpappap (circled at the beginning) divided to generate daughters (circled at the end of video) before ventral enclosure was complete in this embryo. All nuclei are labeled with GFP and membranes of *egl-1*–expressing cells are labeled with mCherry (magenta). Time-lapse images used for this video were obtained using confocal microcopy. Video playback is at 600x real speed. The embryo shown carried the transgenes *stIs10026* and *nIs861*.

**Movie 6 A few cells are extruded from a vehicle-treated MDCK monolayer**

A time-lapse video of mammalian MDCK monolayer treated with vehicle control for 21.25 h shows that a few cells are extruded during this period. Extruded cells can be identified as bright, white, rounded spots rising from the epithelial plane. Video playback is at 7200x real speed. Scale bar, 100 µm.

**Movie 7 A large number of cells are extruded from an HU-treated MDCK monolayer**

A time-lapse video of mammalian MDCK monolayer exposed to HU for 21.25 h shows that many more cells are extruded during this period as a result of HU treatment. Extruded cells can be identified as bright, white, rounded spots rising from the epithelial plane. Video playback is at 7200x real speed. Scale bar, 100 µm.

## KEY RESOURCES TABLE

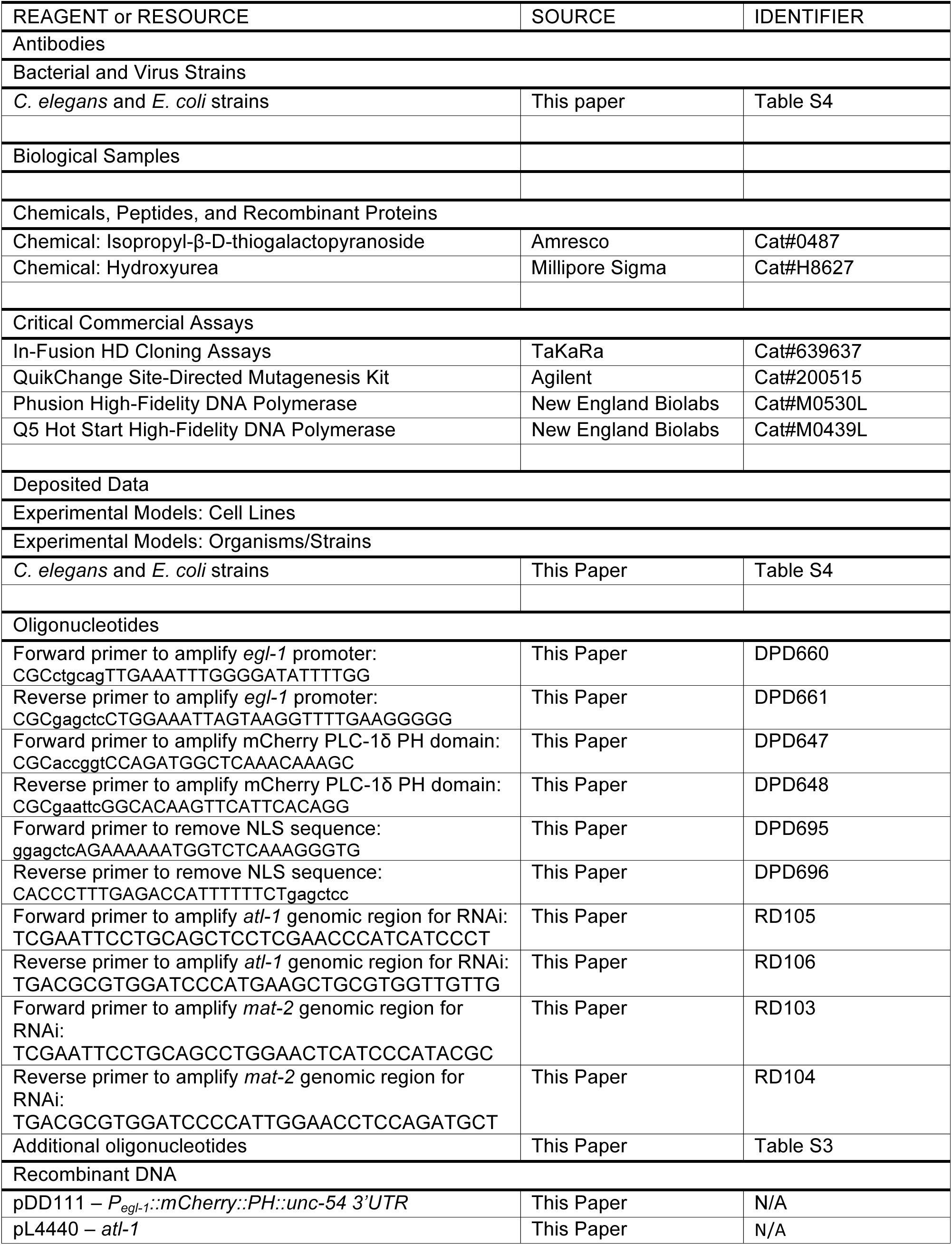
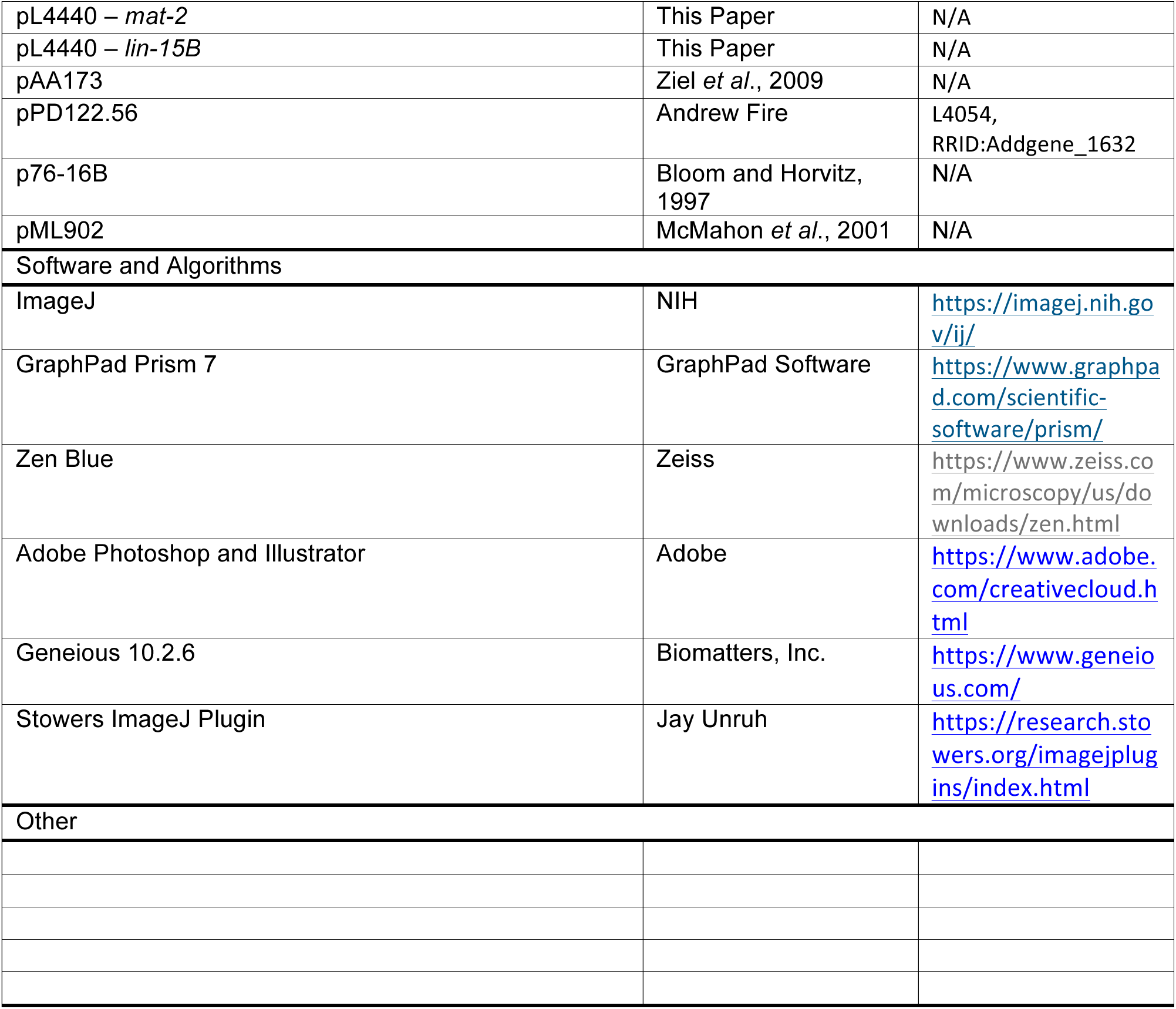

**Table S3.**
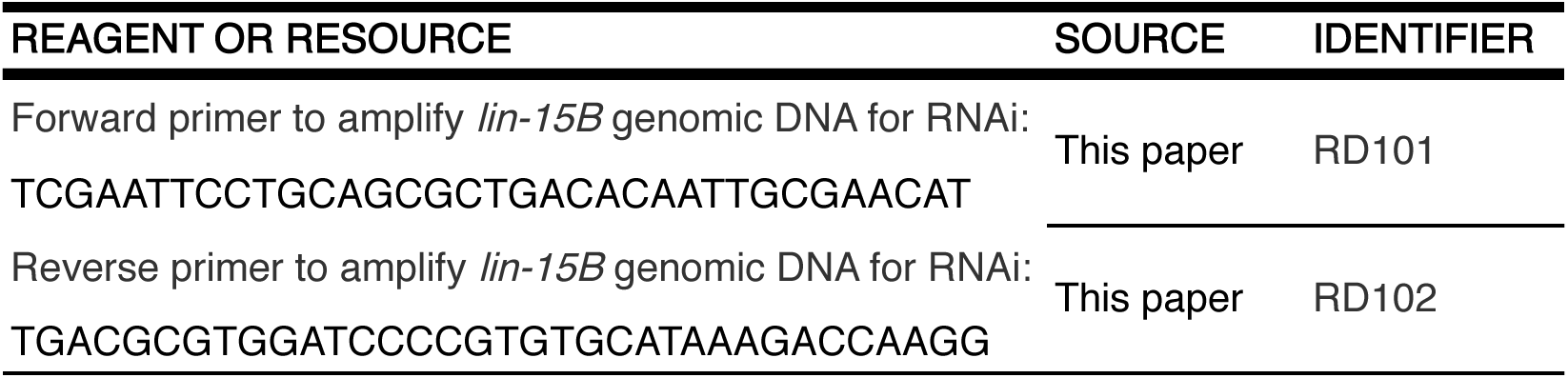
Additional Oligonucleotides.

**Table S4.**
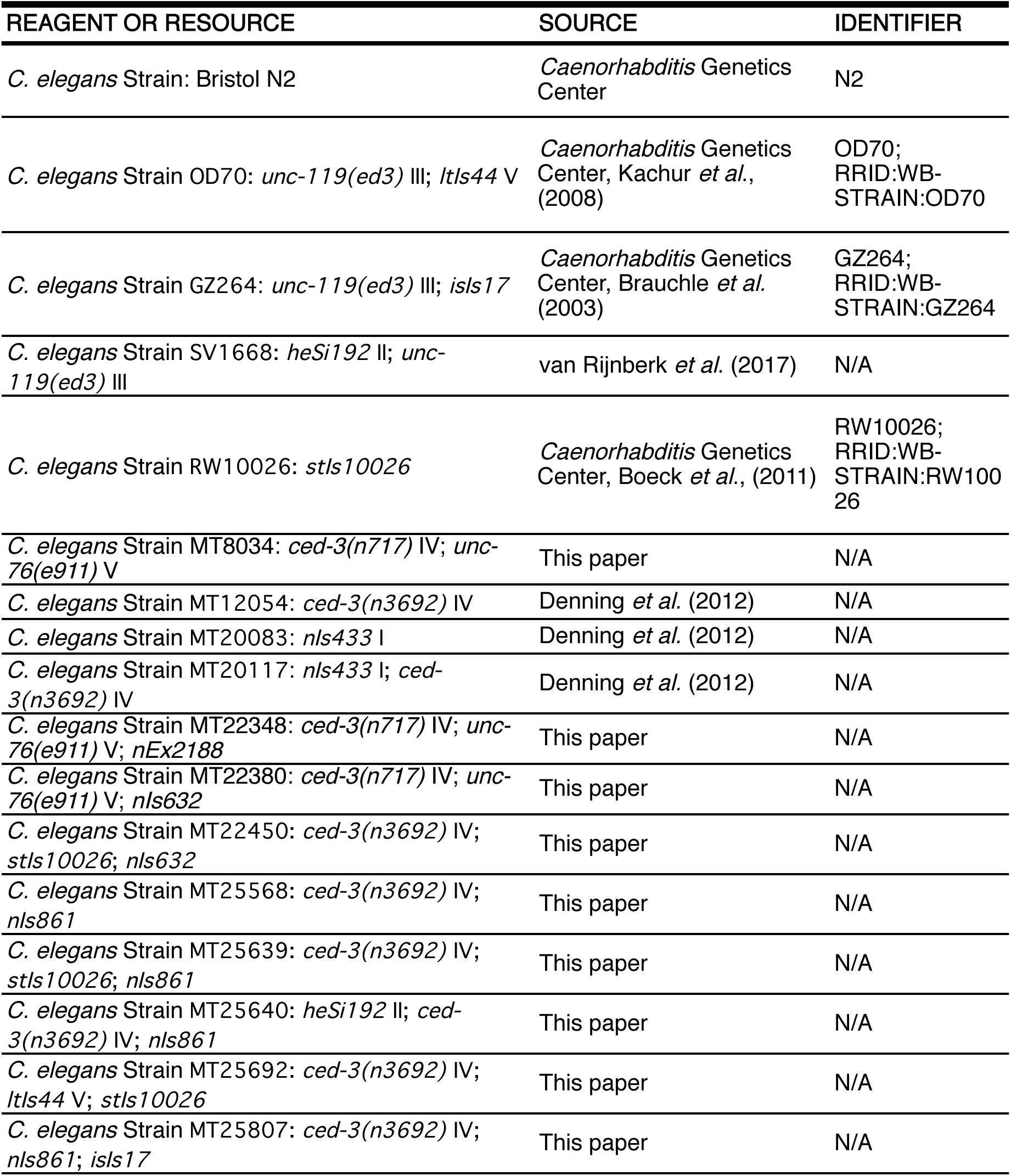
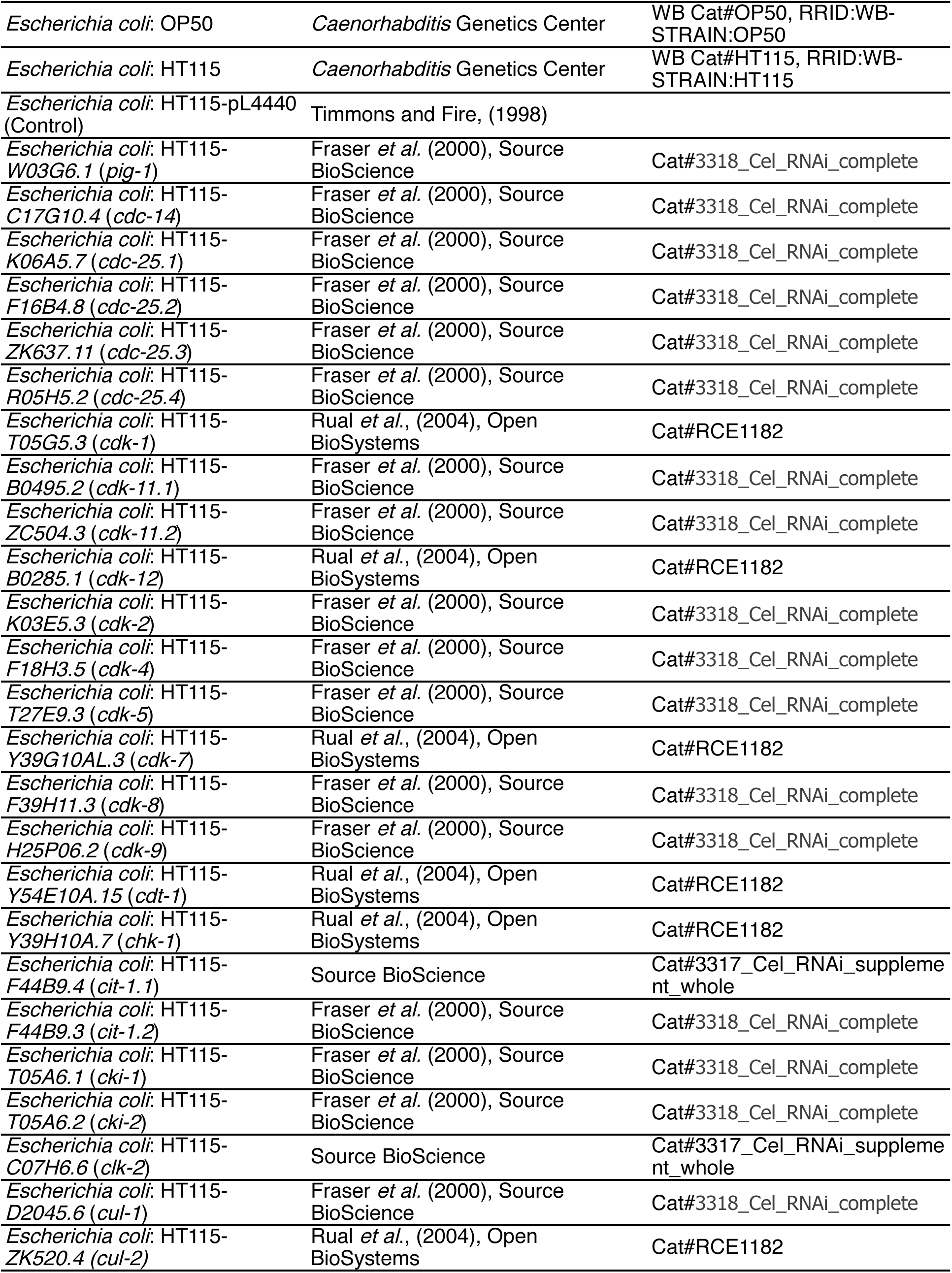
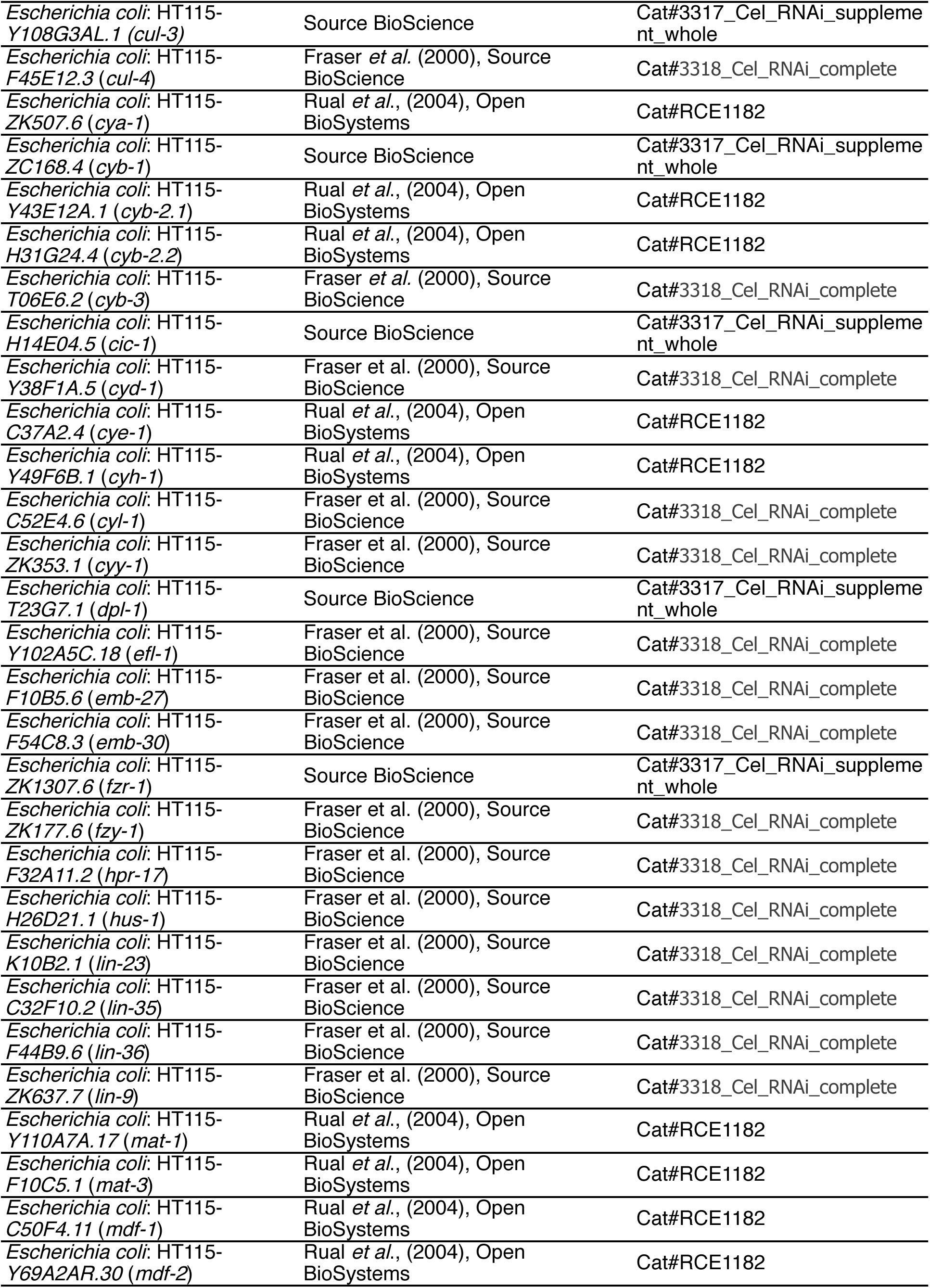
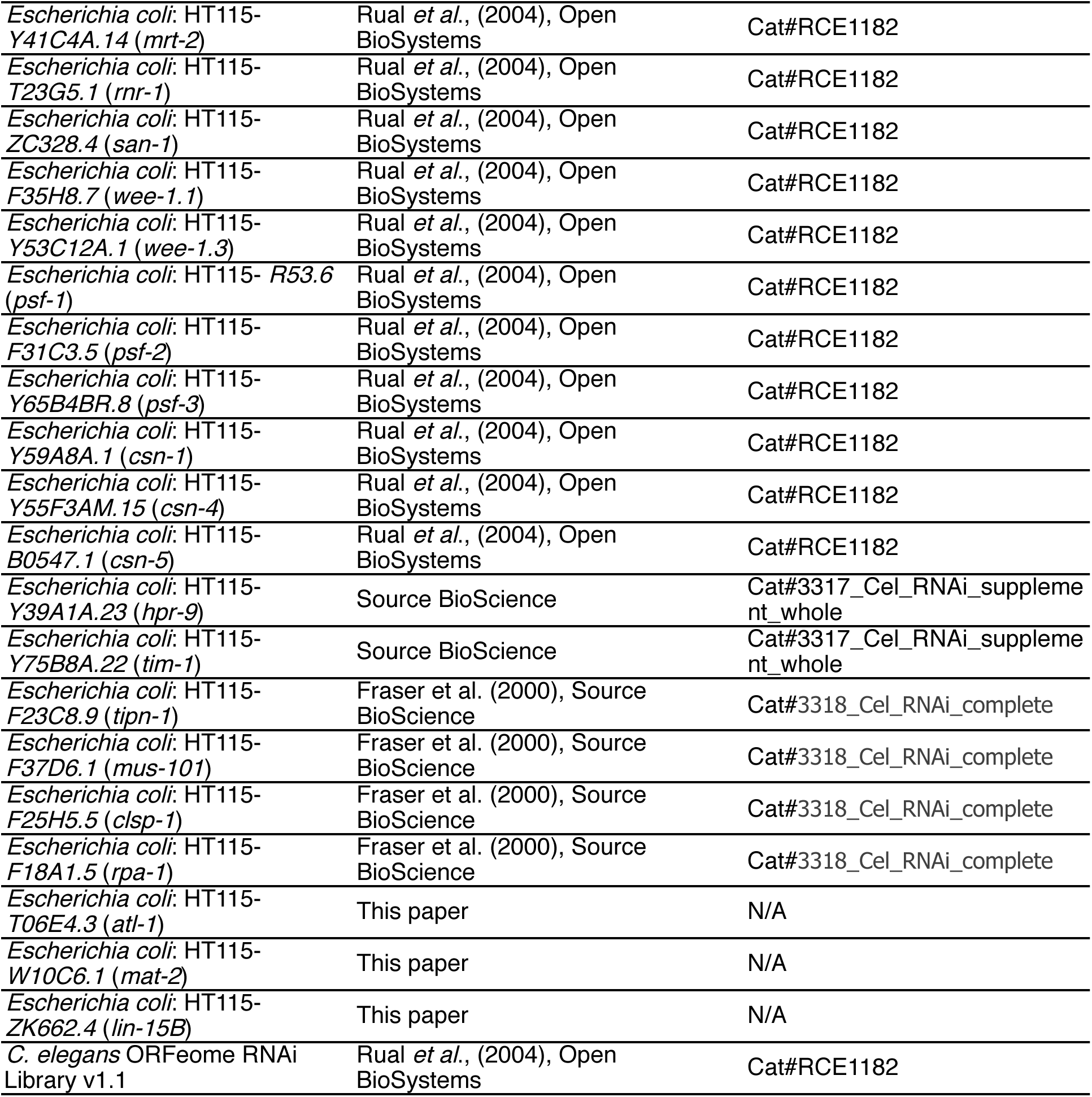
*C. elegans* and *E. coli* strains.

## Materials and Methods

### Plasmids

L4054 was a gift from Andrew Fire (Addgene plasmid # 1632; http://n2t.net/addgene:1632; RRID:Addgene_1632). pDD111 - *P_egl-1_::mCherry::PH::unc-54 3’UTR* was generated with the following steps: i) 6.8 Kb of the *egl-1* promoter was amplified from genomic DNA with Phusion DNA polymerase using the primers DPD660 and DPD661; ii) the amplicon was digested with PstI and SacI (New England Biolabs) and ligated into pPD122.56, which encodes 4xNLS::GFP to generate *P_egl-1_::4xNLS::GFP::unc-54 3’UTR*; iii) mCherry-PH (<u>P</u>leckstrin <u>H</u>omology) sequence was amplified from pAA173 using DPD647 and DPD648 and digested with EcoRI and AgeI (New England Biolabs) and ligated into the pDD122.56 - *P_egl-1_::4xNLS::GFP::unc-54 3’UTR*, which generated the plasmid pDD122.56 - *P_egl-1_::4xNLS::mCherry::PH::unc-54 3’UTR*; iv) the 4xNLS sequence was removed with the primers DPD695 and DPD696 using QuikChange Site-Directed Mutagenesis (Agilent) to generate pDD111 - *P_egl-1_::mCherry::PH::unc-54 3’UTR*.

RNAi clones were constructed for *atl-1*, *mat-2* and *lin-15B*. Genomic regions of about 1 kb were amplified from wild-type genomic lysates using Q5 Hot Start high-fidelity polymerase (New England Biolabs) with the following primers: *atl-1*

RD105 TCGAATTCCTGCAGCTCCTCGAACCCATCATCCCT

RD106 TGACGCGTGGATCCCATGAAGCTGCGTGGTTGTTG

*mat-2*

RD103 TCGAATTCCTGCAGCCTGGAACTCATCCCATACGC

RD104 TGACGCGTGGATCCCCATTGGAACCTCCAGATGCT

*lin-15B*

RD101 TCGAATTCCTGCAGCGCTGACACAATTGCGAACAT

RD102 TGACGCGTGGATCCCCGTGTGCATAAAGACCAAGG

These inserts were cloned into the pL4440 vector linearized with *XmaI* (New England Biolabs) using the In-Fusion HD cloning kit (TaKaRa) according to manufacturers instructions. The cloned vector was then transformed into competent HT115 bacterial cells. Correct RNAi clones were identified by Sanger sequencing. Geneious 10.2.6 (Biomatters, Inc.) was used to guide all plasmid design and construction.

### Contact for Reagent and Resource Sharing

Further information and resource sharing requests should be directed to and will be fulfilled by the lead contact, H. Robert Horvitz.

### Strains, transgenes and mutations

*C. elegans* hermaphrodite strains were maintained on Nematode Growth Medium (NGM) plates containing 3 g/L NaCl, 2.5 g/L peptone and 17 g/L agar supplemented with 1 mM CaCl_2_, 1 mM MgSO_4_, 1 mM KPO_4_ and 5 mg/L Cholesterol with E. coli OP50 as a source of food (Brenner, 1974). All strains were derived from Bristol N2 and are listed in Table S2. *ced-3(lf)* refers to the *n3692* deletion allele of *ced-3* (Denning *et al*., 2012). *C. elegans* strains carrying the transgenes *nIs861* and *isIs17* were maintained at 25°C. All other strains were maintained at 22°C. The transgenes and mutations used are listed below:

**LGI:** *nIs433[P_pgp-12_::4xNLS::GFP::unc-54 3’UTR; p76-16B(unc-76(+))]*
**LGII:** *heSi192[P_eft-3_::tDHB::eGFP::tbb-2 3’UTR + Cbr-unc119(+)]*
**LGIII:** *unc-119(ed3)*
**LGIV:** *ced-3(n3692, n717)*
**LGV:** *unc-76(e911), ltIs44[P_pie-1_::mCherry::PH(PLC1delta1) + unc-119(+)]*
**Unknown linkage:** *stIs10026[P_his-72_::HIS-72::GFP], isIs17[pGZ295(P_pie-1_::GFP::pcn-1(W03D2.4)), pDP#MM051 (unc-119(+))], nIs861[pDD111(P_egl-1_::mCherry::PH::unc-54 3’UTR)], nIs632[pDD111(P_egl-1_::mCherry::PH::unc-54 3’UTR), pML902 (dlg-1::GFP),p76-16B(unc-76(+))]*
**Extrachromosomal array:** *nEx2188[pDD111(P_egl-1_::mCherry::PH::unc-54 3’UTR), pML902 (dlg-1::GFP), unc-76(+)]*

*nIs632* and *nIs861* express membrane-localized mCherry from the *egl-1* promoter, which facilitated the identification of ABplpappap (an *egl-1* expressing cell). *nIs632* does not express *dlg-1::GFP*, presumably as a result of partial transgene silencing (Hsieh *et al*. 1999; Grishok *et al*. 2005; Fischer *et al*. 2013). *stIs10026* (Boeck *et al*., 2011) ubiquitously expresses a GFP-tagged histone HIS-72 from its endogenous promoter, which produces fluorescence in the nuclei of all cells and facilitates in providing the context in which extrusion events are observed.

### Germline transformation

Transgenic lines were generated using the standard germline transformation procedure (Mello *et al*., 1991). Extrachromosomal array transgene *nEx2188* was generated by injecting pML902 at 3 ng/µL, pDD111 at 40 ng/µL, p76-16B (unc-76(+)) at 60 ng/ul and 1Kb Plus DNA ladder (Thermo Fischer Scientific) at 50 ng/µl into *ced-3(n717) IV; unc-76(e911) V* double mutant animals. *nIs632* was generated by gamma-ray irradiation (4,800 rads) of *nEx2188*-carrying L4 animals and was identified by the 100% transmission of the transgene from transformed parent to progeny. *nIs861* was a spontaneous integration in a germline cell of an animal injected with pDD111 at 10 ng/µL and 1 kb DNA ladder at 90 ng/µL, and was identified by the 100% transmission of the transgene from transformed parent to progeny.

### RNAi treatments and genome-wide RNAi screen

Previously described feeding RNAi constructs and reagents were used to perform RNAi feeding experiments (Fraser *et al*., 2000; Rual *et al*., 2004). Briefly, HT115 *Escherichia coli* bacteria carrying RNAi clones in the pL4440 vector were grown for at least 12 h in Luria broth (LB) liquid media with 75 mg/L ampicillin at 37°C. These cultures were seeded onto 6 cm Petri plates with Nematode Growth Medium (NGM) containing 1 mM isopropyl-β-D-thiogalactopyranoside (IPTG) (Amresco) and 75 mg/L ampicillin and incubated for 24 h at 22°C. For imaging experiments using confocal microscopy, 10 L4 animals were added to each RNAi plate and imaging of progeny embryos was performed on the next day as described in Microscopy below. For excretory cell counts, five L4 animals were added to each RNAi plate and L3-L4 progeny were scored for number of excretory cells, as described in Excretory cell count below. In case a bacterial clone targeting a certain gene was not available in previously constructed libraries (Kamath *et al*., 2003; Rual *et al*., 2004), we generated our own RNAi clone as described in Molecular biology above.

The ORFeome RNAi library was used to conduct a genome-wide RNAi screen (Rual *et al*., 2004). For each day of the RNAi screen, all bacterial colonies from two 96-well plates were cultured for at least 12 h at 37°C in LB with 75 mg/L ampicillin. These cultures were then pre-incubated with 1 mM IPTG (Amresco) for 1 h to maximize induction of dsRNA production. 24-well plates with each well containing 2 mL NGM medium with 1 mM IPTG (Amresco) and 75 mg/L ampicillin were prepared in advance and stored at 4°C until needed; they were brought to room temperature a few hours before seeding. Each bacterial colony culture was then seeded onto an individual well of a 24-well plate and incubated for 24 h at 20°C. Three L4 animals were picked into a 10 µl drop of M9 medium, which facilitated their transfer into a well using a pipette. The progeny of these 3 animals were screened 3 days later. Each set of RNAi clones screened also included a *pig-1* RNAi positive control and an empty pL4440 vector negative control. The scorer was blinded to the identity of the RNAi clones. Excretory cell counts were performed as described in Excretory cell counts below. Sanger sequencing was used to confirm the identity of RNAi clones that reproducibly generated a Tex phenotype for more than 10% of the animals scored.

### Microscopy

All RNAi screens scoring excretory cells were performed using a Nikon SMZ18 fluorescent dissecting microscope. DIC and epifluorescence images were obtained using a 63x objective lens (Zeiss) on an AxioImager Z2 (Zeiss) compound microscope and Zen Blue software (Zeiss).

For confocal microscopy, embryos staged at the 200-300-cell stage were picked and mounted onto a glass slide (Corning) with a freshly prepared 2% agarose pad. Embryos with ventral surfaces facing the objective were selected for imaging. Confocal images were obtained using a 63x objective lens (Zeiss) on a Zeiss LSM800 confocal microscope.

For observing extrusion (or absence of extrusion), we focused particularly on the cell ABplpappap, the identification of which is facilitated by its central position on the ventral surface (Sulston *et al*., 1983). The fluorescent transgene *nIs861[Pegl-1::mCherry::PH]* or *nIs632[Pegl-1::mCherry::PH; dlg-1::GFP]*, which express the Pleckstrin homology domain of PLC-δ fused to mCherry from the promoter of *egl-1*, was used to label the membrane of the ABplpappap cell, an *egl-1* expressing cell (Denning *et al*., 2012), to further facilitate cell identification. Another fluorescent transgene *stIs10026[his-72::GFP]*, which expresses GFP-tagged HIS-72 histone protein, was used to label the nuclei of all cells to help define ABplpappap’s location within the embryo. Time-lapse confocal microscopy was used to monitor the location of ABplpappap in embryos, keeping the cell in view by refocusing on it every 30 sec. Confocal imaging during a period of about 50 min during which ventral enclosure (migration and meeting of hypodermal cells on the ventral surface of the embryo) occurs was sufficient to determine whether ABplpappap did or did not undergo extrusion.

For determining whether ABplpappap and other cells that are extruded entered the cell cycle, the transgene *heSi192[Peft-3::tDHB::eGFP::tbb-2 3’UTR]* was used to express a codon-optimized (for *C. elegans*) C-terminal fragment of Human DNA Helicase B, which translocates from the nucleus to the cytoplasm in response to the activity of the cell cycle CDKs 1 and 2 (van Rijnberk *et al*., 2017). *nIs861* was used to label the membrane of ABplpappap with mCherry to facilitate cell identification.

For determining the cell cycle phase of ABplpappap and other extruded cells, *isIs17[Ppie-1::GFP::PCN-1]* was used to express GFP-tagged PCN-1 protein, which produces a phase-specific fluorescence intensity and localization pattern. *nIs861* was used to label the membrane of ABplpappap with mCherry to facilitate cell identification.

Images were processed with ImageJ software (NIH), Photoshop CC 2019 (Adobe) and Illustrator CC 2019 (Adobe) software. The Time Stamper function in the Stowers ImageJ plugin was used to mark elapsed time on time-lapse videos.

### Excretory cell counts

Excretory cell counts were performed using a dissecting microscope equipped with fluorescence at a total magnification of 270x. For the genome-wide RNAi screen, roughly 50 animals were examined in each well of a 24-well plate and any well with more than 5 animals with two excretory cells was marked for confirmatory testing. Excretory cell counts in confirmatory RNAi experiments, candidate RNAi experiments and experiments with genetic mutants were conducted using 6 cm Petri plates with appropriate media. Animals were first immobilized by keeping the Petri plates on ice for 30 min. At least 100 animals at the L3-L4 larval stage were scored for each genotype or RNAi experiment unless there was extensive lethality or a growth defect, in which case a lower number or earlier-stage animals, respectively, were scored. A cell was scored as an excretory cell if it was located in the anterior half of the animal and its nucleus had strong GFP expression.

### tDHB-GFP fluorescence intensity quantification

The ABplpappap nuclear boundary, cell membrane boundary and the tDHB-GFP fluorescence signal were determined from DIC, mCherry and GFP channels, respectively, of confocal images of RNAi treated *ced-3(lf)* embryos expressing the transgenes *heSi192* and *nIs861*. Mean tDHB-GFP fluorescence intensities inside the nuclear region, entire cell and background were quantified using Fiji software. Mean cytoplasmic tDHB-GFP fluorescence intensity was calculated by the following formula

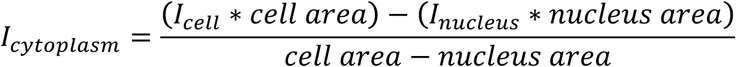

*I_cytoplasm_*, *I_cell_* and *I_nucleus_* denote the mean tDHB fluorescence intensity in the cytoplasm, cell and nucleus, respectively. The ratio of nuclear-to-cytoplasmic tDHB fluorescence intensity in Figure 3H was adjusted for background fluorescence (measured from a random area outside the embryo boundaries), i.e., the background fluorescence intensity was subtracted from both nuclear and cytoplasmic fluorescence intensity values before calculating the ratios.

### Calculation of cell size

Confocal micrographs were obtained for multiple focal planes starting at the ventral surface and ending at the dorsal surface of the embryo, with each plane separated by a distance of 0.37 µm. The greatest area occupied by a cell in any plane was designated the “maximum area” of a cell.

### Cell culture

MDCK and MDCK-Fucci (Streichan et al., 2014) cells were cultured in DMEM supplemented with 10% fetal bovine serum and 1% penicillin/streptomycin in a humidified incubator at 37°C with 5% CO_2_.

### Chemicals

2 mM HU (Millipore Sigma, Cat#H8627) was prepared in culture medium prior to each experiment.

### Mammalian cell imaging

These assays were performed using 6-well plastic plates. 20,000 MDCK cells were seeded in each well and grown to confluence for 72 h. The day of the experiment, cells were washed twice with PBS and treated with fresh medium or 2 mM HU in medium. After equilibration, plates were imaged at 15-min intervals for up to 24 h, using an Evos M7000 imaging system equipped with a humidified onstage incubator (37°C, 5% CO_2_). Several positions per well were imaged in the phase contrast and green and red fluorescence channels available in this system.

### Mammalian cell extrusion quantification

In time-lapse phase contrast images, extruding cells are easily identifiable as bright, white, rounded spots emerging from the epithelial plane. We counted the number of cells with these features for each condition using the Cell Counter plugin of Fiji (Schindelin *et al*., 2012). Extrusions are reported as *number of extruding cells/h* for comparison between experiments of different duration.

### Mammalian cell cycle phase determination

The Fucci system differentially labels the nuclei of cells in G1 (red) and S/G2/M (green) (Sakaue-Sawano *et al.*, 2008). Images of MDCK-Fucci cells with HU or control treatment were obtained in the phase contrast, red and green fluorescence channels as per Mammalian cell imaging above. For each position, a multi-channel stack was built using Fiji (Schindelin *et al*., 2012). After identifying an extruded cell in the phase contrast channel, the cell cycle phase was determined using the fluorescence channels.

### Mammalian re-seeding experiments

At the end of an imaging experiment, supernatants were collected and centrifuged (1200 rpm, 5 min, room temperature). Pellets were re-suspended in 50 µL of PBS, and 10 µL of the suspension was used for cell counting with Trypan blue in a Neubauer chamber, allowing us to simultaneously calculate the number of cells being re-seeded and the fraction of cells that was apoptotic. The remaining cells were seeded with 1 mL of fresh medium in a 24-well plate and grown in the cell culture incubator. Pictures were taken at 2 h and 24 h for cell counting.

### Statistical analysis

For calculation of statistical significance for ratios, the ratios were first transformed to logarithm values. Ordinary one-way ANOVA was performed to determine statistical significance of the ratios with the assumption that logarithm of ratios produced a normal distribution of values. The maximum area of ABplpappap was also assumed to have normal distribution under different RNAi conditions and ordinary one-way ANOVA was used to determine statistical significance. Normal distributions with unequal variances were assumed for rates of extrusion under HU and vehicle treatments, and Welch’s two-tailed t-test was performed to determine statistical significance. No assumptions were made about the distributions of the rates of apoptosis under HU and vehicle treatments, and hence the Mann-Whitney test was used to determine statistical significance. No assumptions were made about the distribution of fraction of extruded cells in different phases of the cell cycle after HU and vehicle treatments, and the Kruskal-Wallis test was used to determine statistical significance. Normal distribution was assumed for numbers of cells reseeded in fresh media after pre-treatment in different conditions, and ordinary one-way ANOVA was used to determine statistical significance. All statistical analysis was performed using Prism 7 (GraphPad Software).

